# Tissue mechanics and somatosensory neural responses govern touch sensation in *C. elegans*

**DOI:** 10.1101/471904

**Authors:** A. Sanzeni, S. Katta, B.C. Petzold, B.L. Pruitt, M.B. Goodman, M. Vergassola

## Abstract

The sense of touch hinges on tissues transducing stimuli applied to the skin and somatosensory neurons converting mechanical inputs into currents. Like mammalian Pacinian corpuscles, the light-touch response of the prime model organism *C. elegans* adapts rapidly, and is symmetrically activated by the onset and offset of a step indentation. Here, we propose a quantitative model that combines transduction of stimuli across the skin and subsequent gating of mechanoelectrical channels. For mechanics, we use an elastic model based on geometrically-nonlinear deformations of a pressurized cylindrical shell. For gating, we build upon consequences of the dermal layer’s thinness and tangential stimuli. Our model demonstrates how the onset-offset symmetry arises from the coupling of mechanics and adaptation, and accounts for experimental neural responses to a broad variety of stimuli. Predicted effects of modifications in the mechanics or the internal pressure of the body are tested against mechanical and neurophysiological experiments.

## INTRODUCTION

The sense of touch is a prime example of mechanotransduction in biology [1–4]. Mechanical stimuli applied to the skin are shaped by tissue properties and converted by somatosensory neurons into currents, which ultimately control perception and behavioral responses. A notable instance is the response of the nematode *C. elegans* to light touches of its head, which make the animal rapidly back away from the stimulus, and suppress its head movements [5, 6]. The ethological importance of this reaction is arguably associated with escape from predacious fungi that cohabit with nematodes in organic debris [7]. The escape phenotype, along with the ability to obtain whole-cell patch-clamp recordings from identified mechanoreceptor neurons in living animals, make *C. elegans* ideally suited to advance understanding of mechanosensation [8, 9]. Major open questions are how external mechanical loads activate sensory mechanoelectrical transduction (MeT) channels, and how the skin shapes mechanical cues delivered by touch stimuli [10].

Ion channels along Touch Receptor Neurons (TRNs) are gated mechanically and rapidly adapt to a constant stimulus [11]. Similar properties are observed in other mechano-sensitive channels, e.g. channels of large and small conductance that control the osmotic pressure in bacteria [12, 13], as well as the channels responsible for vertebrate hearing [14, 15]. Unlike those cases, TRNs and other *C. elegans* mechanoreceptor neurons [16] show a similar response when an external stimulus is applied or released. A rapidly-adapting, symmetric response is also observed in mammalian Pacinian corpuscles, where it is due to the onion-like lamellar capsule that encases its nerve ending [17, 18]. Since this peculiar structure is not observed in most other receptors, such as *C. elegans* TRNs, an alternative explanation is required.

In [19], we introduced a possible mechanism that could account for the response of TRNs and its symmetry. In its simplest implementation, an elastic filament connected to the channel is stretched at the onset of the stimulus, which results in a gating force on the channel that modulates its opening probability. Adaptation results from the relaxation of the filament, with a timescale controlled by the balance between elastic and frictional forces. At the offset of the stimulus, the elastic link is stretched in the opposite direction with respect to the onset, which would provide similar gating forces and response. The model was inspired by the tip-link model for hair cells [14, 15] yet it differs in that the tangential, and not the vertical, stretching of the filament is relevant. Pictorially, the gating of hair cells’ channels by a hinged trapdoor was replaced by a sliding trapdoor. The importance of tangential stimuli relates to the thinness of the tissue where TRNs are embedded, and to the mechanics of thin shells [20–22].

While appealing in its simplicity, the model in [19] is incomplete: the transfer of stimuli across the skin was not accounted, and channels along the TRNs were replaced by a single effective one, to bypass the spatial dependence of the deformation field. The proposal in [19] calls for a stronger foundation and validation, as well as a physical connection with tissue mechanics. That is our goal here.

Beyond methodological interest, a complete model that encompasses tissue mechanics and channel activation, allows us to address a number of open issues: the role played by *C. elegans* body mechanics and internal pressure (as well as their mutations) on touch response; the response to general patterns of stimulation; how the onset/offset symmetry of the touch response relates to the transduction of stimuli, and whether the symmetry holds for single channels or at the population level only. Here, we analyze mechanics in the regime of large deformations, combine it with the modeling of channel activation, and thereby obtain experimentally testable predictions on the mechanical and neural response to touch. Their validity will be supported both by previous and new experimental data.

## RESULTS (MODELING)

The following Sections reflect the two elements that govern the sense of touch: the mechanics of the nematode body and the gating of the channels along TRNs.

### *C. elegans* body mechanics

The body of *C. elegans* hermaphrodites consists of an outer and an inner tube, separated by a fluid-filled pseudocoelom. The former comprises cuticle, hypodermis, excretory system, neurons, and longitudinal muscles; the latter is formed by pharynx, intestine, and gonad [23]. Following previous work [24], this body structure suggests the model of a cylindrical elastic shell with internal pressure (see Fig. 1), which accounts for observations that *C. elegans* burst when their cuticle is punctured, as first reported for the related nematode *Ascaris* [25]. Internal pressure subsumes the combined effects of internal organs and the fluid-filled pseudocoelom. Elasticity of the external shell provides an effective description (see conclusions for viscoelastic effects) of the mechanics in the outer tube.

Natural stimuli substantially indent the body of a worm; typical experiments feature a bead of radius 5 − 10*μm* indenting the body to a maximum depth of ∼ 10*μm*. That is about half the radius of the shell, and is larger than its thickness ∼ 1*μm*.

#### The mechanical elastic model

The simplest physical model consistent with the above observations has that the strain within the shell is modest yet the displacement of material points can be substantial. The latter implies that the linear approximation of the strain is not appropriate and must be replaced by the nonlinear Green-Lagrange expression:

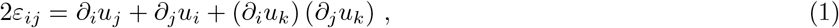

where ∂_j_ is the derivative with respect to the spatial coordinate *x_j_* (*j* = 1, 2, 3) and *u_i_* is the *i*-th component of the displacement. The equations for the stress tensor *σ_ij_* (Piola-Kirchoff of the second type) read [20–22]

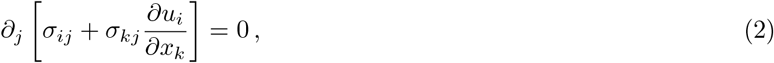

where sum over repeated indices is implied. Quadratic terms in Eqs. (1) and (2) make them appropriate for large deformations. Nonlinearities are usually called “geometric” due to their relation to the shape of material elements [21, 22]. Linear approximations that neglect geometric nonlinearities can lead to substantial discrepancies, as shown below.

The linear Hookean relation between stress and strain 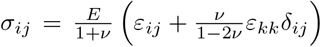 is consistent even for large deformations provided gradients of displacement stay moderate, which will be the case here (see Appendix A). Here, *E* is Young’s modulus, *ν* is the Poisson ratio, and *σ_ij_* is energy conjugate to *ε_ij_*, i.e. the elastic energy *Ε_el_* upon a deformation varies as *δΕ_el_* = *∫ σ_ij_ δε_ij_ dV*.

In addition to *E* and *ν*, parameters of the model are (see Fig. 1): the length *L* and the thickness *t* of the shell, the radius *R* of its middle surface, and the internal pressure *p*. The pressure *p* should be understood as the difference between the internal and the external atmospheric pressure, which has minor effects for our thin cylindrical shells (see Appendices B and C) and we shall neglect here to simplify the following dimensional analyses. Elastic parameters are effective quantities that subsume different contributions in the inner and outer tube. Previous estimates of those parameters are discussed in the Section on Results for experimental validations. Boundary conditions generated by the pressure *p* and the external forces are in the next Section. Finally, Appendix D discusses the reduction of the 3D Eqs. (2) to the 2D thin-shell limit.

**FIG. 1.**
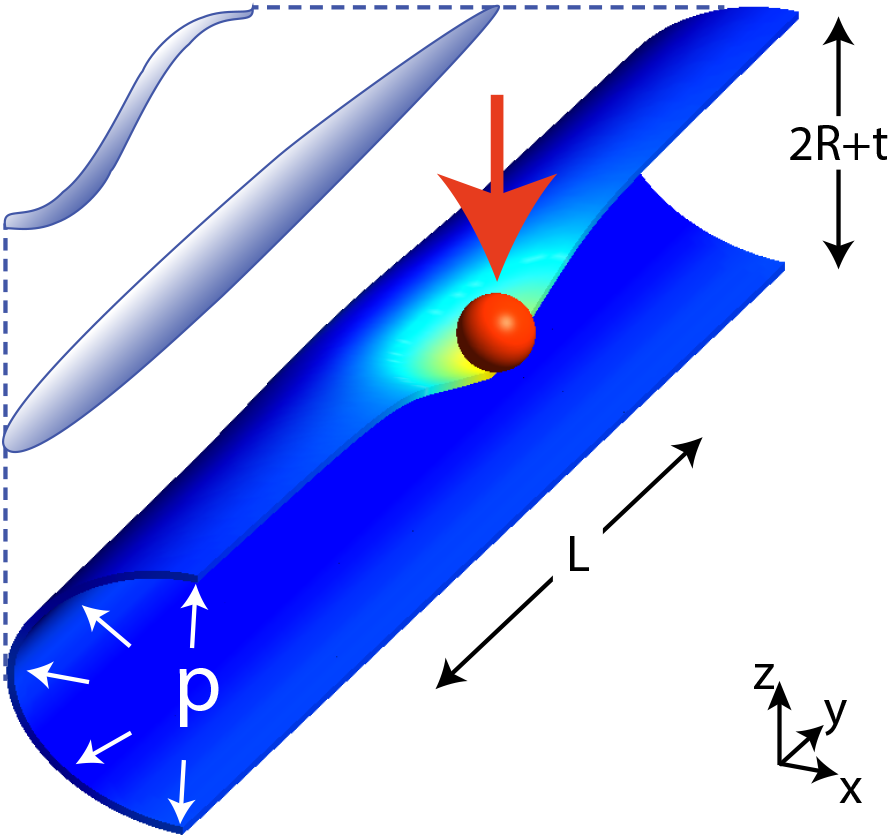
Scheme of the geometry in our model for *C. elegans* mechanics. The figure shows the scheme of a worm in a natural posture (left), straightened (as in neurophysiology experiments), and the model (right) that we shall consider here: a cylinder of length *L* ≃ 1 *mm* and radius *R* + *t*/2 ≃ 25*μm* is indented by a spherical bead (with radius 10*μm* unless stated otherwise), applied here at its center. *R* is the radius of the middle surface and *t* is the thickness of the shell. Only half of the cylinder is shown for clarity.

#### Numerical simulations of the elastic model

The nonlinear structure of Eq. (2) hampers analytical approaches, which pushed us to numerical finite-element methods (see [26] for an introduction). Numerical simulations of Eq. (2) were performed by the open-source program *code-aster* [27]. An hexahedral element with 8 standard nodes (HEXA8) was used in combination with a mesh sensitivity analysis to verify that results are minimally sensitive to the element size. The numerical procedure was benchmarked and tested by comparing its results to known elasticity problems. In particular, Appendix E reports on the comparison with the deformation field and the force-indentation relation produced in small indentations of cylindrical shells, where an analytical solution is available [28], as well as large indentations of pressurized spherical shells, where a simplified equation was derived in [29]. In all cases, agreement was verified.

For the internal pressure *p*, we shall make the simplest possible hypothesis that *p* holds constant when the stimulus is exerted onto the body of the worm. While active readjustments of the internal pressure could *a priori* be accommodated in our approach, results reported below indicate that the simplest option is sufficient.

As for boundary conditions, neurophysiology experiments have worms glued onto a plate and limited in the vertical displacement of their body’s lower half (see Materials and Methods). Since a mathematical formulation is not obvious (and elasticity has long-range effects), we tested different boundary conditions and present two of them for comparison (see Appendix F). For the first, the lower half of the body is vertically rigid, *i.e*. the upper half of the cylinder in Fig. 1 is free to move, while the lower one is constrained to move only parallel to the plane onto which the worm is glued. For the second boundary condition, the lower half is fixed, *i.e*. allowed to move neither vertically nor laterally. Results presented in the main text are obtained with the latter condition. As for the two ends of the cylinder, we present results with vanishing lateral forces; Appendix G discusses the effects of plugs at the ends.

### The channel gating mechanism

In this Section we address the activation of MeT channels by mechanical stimuli. Our presentation recapitulates and extends the model in [19]. TRNs extend below the cuticle for about half the longitudinal length of the nematode. Channels along the TRN [30] are stimulated by the deformations described in the previous Section. Individual channels contribute to the total TRN current, which is the quantity measured in experiments that we seek to account for and predict.

#### Stimulation and adaptation of a channel

The mechanism in [19] posits that the dynamics of individual channels is the combination of an elastic and a relaxation (frictional) component. While various implementations may be contemplated, we shall refer for concreteness to a situation where each ion channel is connected to an elastic filament. We denote by **r**^*c,f*^ the undeformed positions of the channel and the tip of its elastic filament; the corresponding displacements induced by the deformation of the embedding tissue are **Δr**^*c,f*^.

The elastic component reflects the Hookean response of the filament to its stretching. Elastic energy is 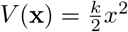, where x = **Δr**^f^ − **Δr**^c^ is the elongation of the filament with respect to its undeformed configuration. The corresponding restoring force is

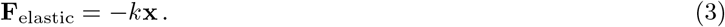

As for the frictional component, the TRN and its channels are embedded in the medium and expected to move with it, i.e. **Δr**^*c*^ = **u**(**r**^*c*^) where *u* is the displacement in (2). Conversely, as the filament slides with respect to the medium, the friction force is

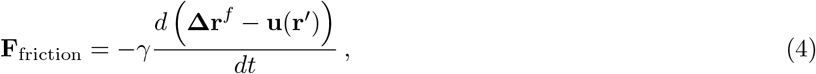

where *γ* is the friction coefficient and **r**′ is the undeformed position of the material point that coincides with the location of the tip, i.e. **r**^*f*^ + **Δr**^*f*^ = **r**’ + **u**(**r**′). Expanding **u**(**r**′) and using that gradients of **u** are small, we obtain **r**’ − **r**^*f*^ ≃ **Δr**^*f*^ − **u** (**r**^*f*^), which is then inserted into (4) to show that **u**(**r**′) can be replaced by **u**(**r**^*f*^). We remind that small gradients of **u** also entered the Hookean relation between stress and strain in the mechanical elastic model.

Effects of inertia are negligible and the overdamped approximation holds at microscopic scales [31], i.e. the sum of the forces **F**_friction_ + **F**_elastic_ = 0, which yields

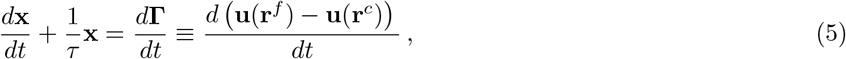

where *τ* = *γ*/*k* is the relaxation time, x is the extension of the filament, and **Δr**^*c*^ = **u**(**r**^*c*^) was used. Eq. (5) drives x to zero for **Γ** constant, which is the basis for adaptation. Eq. (5) is supplemented by the constraints exerted by the neural membrane around the channel, which limit the motion in the vertical direction. The constraint can be written as x · **ŵ**_3_ ≥ 0, where, for every channel, **ŵ**_1,2_ span the plane locally tangential to the neural membrane while **ŵ**_3_ indicates the orthogonal direction. Specifically (see also Appendix H), we define an orthonormal basis 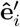 as follows: 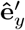 is aligned with the local direction of the (deformed) axis of the cylinder running head-to-tail; 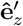 is orthogonal to the neural membrane at the top of the TRN, and oriented outward; 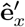 is tangential to the neural membrane, along the remaining direction of a right-handed system. The bases **ŵ**_i_ are constructed by rotating the 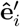 appropriately. For a channel placed at the top of the TRN, the local basis **ŵ**_i_ coincides with 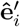. If the channel is rotated by *θ* along the surface of the TRN, then 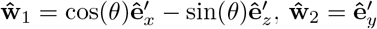 and 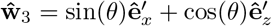.

The dynamics of **Γ** in (5) is obtained by the displacements *u* calculated using the mechanical model. The relation between stretching and *ε* is [20, 22]: Γ^2^(*t*) − Γ^2^(0) = 2*ε_ij_*(*t*)Γ_i_(0)Γ_j_(0), where the Γ’s are again assumed to be small.

#### The opening/closing dynamics of channels

Channels can be in multiple states: open, closed and several sub-conducting [32]. To reduce the number of free parameters, we include a single sub-conducting in between the open and closed states (*C* ⇌ *S* ⇌ *O*). The respective probabilities *P_c,s,o_* obey the master equation

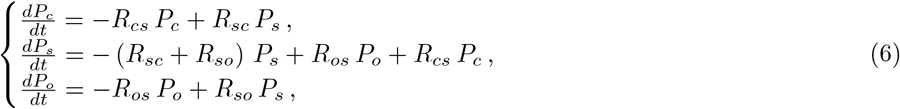

where *R_ij_* are the respective transition rates, and *R_c,o_* = *R_o,c_* = 0 again to minimize free parameters. The channels are posited to work at equilibrium, so that

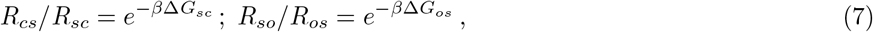

where Δ*G_ij_* is the free energy difference between the states *i* and *j, β* = 1/*k_B_T, T* is the temperature, and *k_B_* is the Boltzmann constant (see, e.g., Ref. [31]).

Channels are coupled to mechanics via their elastic filaments described by Eq. (5). Namely, the extension of the filament modulates the free energy differences among the above states of the channels:

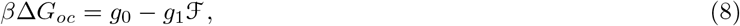

where *g*_0_, *g*_1_ are dimensional constants, and ℱ is the amplitude of the tangential component of **F**_elastic_ in Eq. (3):

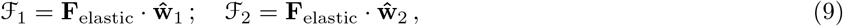

where, for every channel, **ŵ**_1,2_ span the plane locally tangential to the neural membrane while **ŵ**_3_ indicates the orthogonal direction, as defined above.

Choices for the free energy other than (8), e.g. a quadratic dependence on *x*, are discussed in Appendix I. The free energy of the intermediate subconductance state has *a priori* its own parameters. However, to reduce free parameters, its free energy is assumed intermediate between the closed and the open state in Eq. (8)

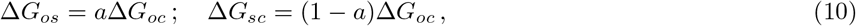

with the only additional parameter 0 ≤ a ≤ 1.

#### The total current flowing along the TRN

Ion channels are believed to be distributed in spots (“puncta”) along the neural membrane [11]. Their distribution is consistent with uniformity in the angular and longitudinal directions, while spacings between successive puncta are distributed log-normally [30]. For simplicity, each punctum is assumed to contain a single channel.

The current along the TRN is the sum *I* = ∑_*k*_ *i_k_* of the currents of individual channels. Its mean is given by

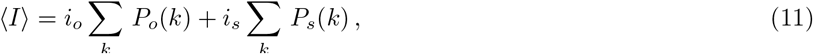

and its variance is calculated similarly (see Appendix J). Here, *i_o_* and *i_s_* are the channel current in its open/subconducting state. *p*(*k*) obey (6), and their rates depends on the position along the TRN via Eqs. (3), (5), (8). The single-channel current *i*_0_ = −1.6 ± 0.2pA, as measured in [11, 32]; other parameters will be inferred from experimental data.

## RESULTS (EXPERIMENTAL VALIDATIONS)

### Experimental validation of the mechanical model

The mechanical model introduced previously will now be tested against existing experimental results on *C. elegans* mechanics [24, 36, 37]. The upshot is that the model captures both indentation data and the measured bulk modulus. We also use the model to infer that stiffness is pressure-dominated rather than shell-dominated, as proposed in [24]. Finally, Appendix K shows that *lon-2* mutants, which alter the cuticle, modify the bulk modulus, contrary to suggestions in [37].

The first step for a proper comparison with experiments is an appropriate choice of the no-stress state: the corresponding length, thickness and radius should be such that the pressurization of the cylinder leads to the values relevant for experiments. In particular, if we want to keep the final (at pressure *p*) values fixed, the no-stress initial values should change as *p* varies. This point, as well as our below results, differ from Ref. [38], where an elastic model is discussed, yet the role of the no-stress state and pressure are not considered. Initial (no-stress) values are conveniently obtained by using perturbative analytical formulæ in Appendix B, which give the variation of various quantities with *p*.

The schematic of an indented shell in our numerical simulations is shown in Fig. 1. Note that the size of the indenter is not negligible with respect to other dimensions, and the region of contact with the cylinder is expected to change with the indentation depth [39].

The thickness of the shell *t* can be rescaled out, as discussed below in the analysis of bending and stretching contributions, and all geometric parameters appearing in Eq. (2) are fixed. The variable factors are *p* and *E*, which can enter the deformation for a given indentation only via their non-dimensional ratio *p*/*E*. We plot then in Fig. 2B the dependence of the deformation profile along the longitudinal coordinate *υs p*/*E*. Vertical deformation is strongest at the center of the indenting bead, and its longitudinal extension decreases with *p*/*E*. The best least squares fit yields *p*/*E* = 0.01.

Fig. 2C shows how the estimate *p* ≃ 40kPa for the internal pressure was obtained: we fix the ratio *p*/*E* = 0.01, predict the relation force-indentation as *p* varies, and make a best fit to the experimental data. The value *p* ≃ 40kPa is consistent with the range 2-30 kPa estimated in the nematode *Ascaris lumbricoides* [25].

The two above estimates yield for the Young’s modulus *E* ∼ 4MPa, which is comparable to the 1.3MPa obtained by measuring the bending stiffness of the nematode [36]. These values differ from prior estimates of 380MPa [24] relied upon formulæ of linear elasticity (valid for indentations ≪ *t*), which do not apply here.

**FIG. 2.**
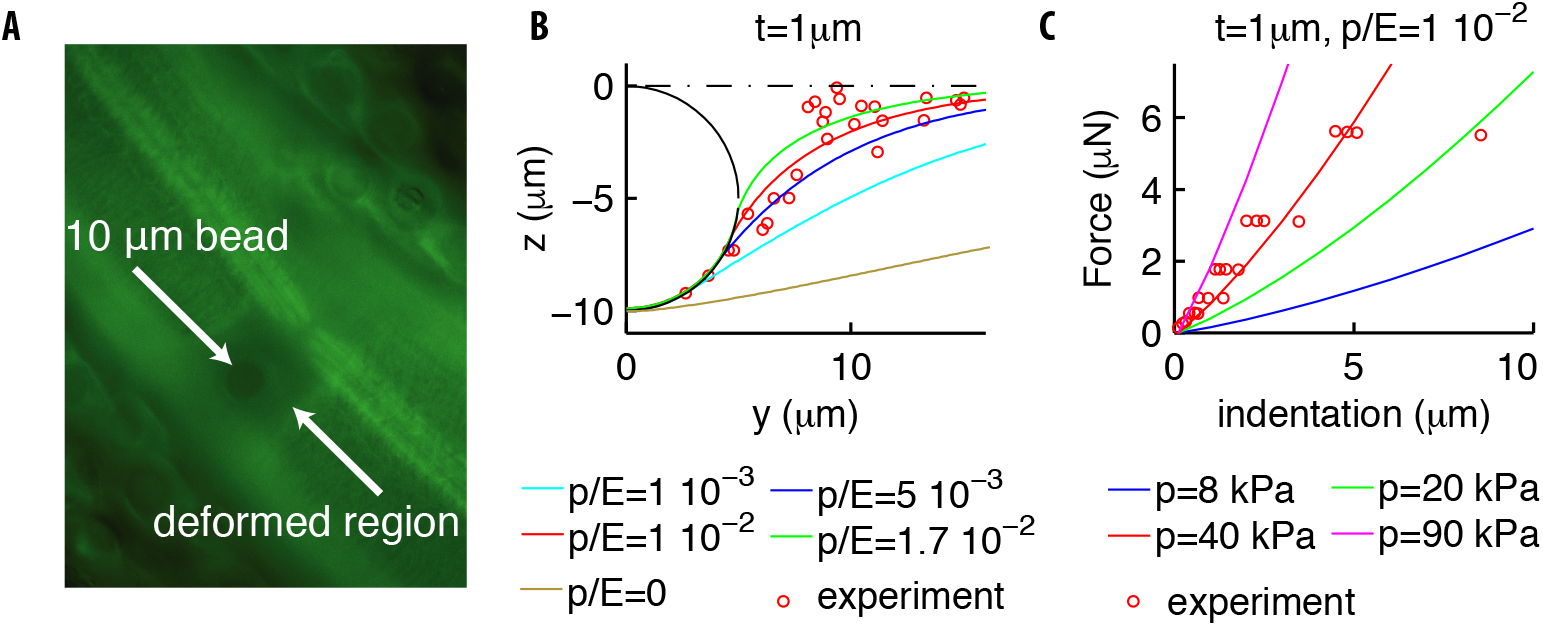
Deformation profiles and force indentation relations. (**A**) Representative photomicrograph of a transgenic animal with GFP-tagged cuticular annuli being pressed into a glass bead. Experimental deformation profiles in (B) were derived from a stack of images at different focal planes. (**B**) Experimental and numerical deformation profiles along the longitudinal axis (the generatrix of the cylinder). Data were obtained as described in Materials and Methods by using 2 biological replicates (adult animals). (**C**) Experimental (from Ref. [19]) and numerical force-indentation relationships. Length is *L* = 1*mm* and the Poisson coefficient *ν* = 0.3.

Having fixed the parameters of our model, we can now independently test it against data on the mechanical response of *C. elegans* to changes in the external pressure [37]. The variation Δ*V* of the initial volume *V*_0_ was found to depend linearly on the variation of the external pressure Δ*p*, and the resulting bulk modulus 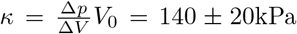. Performing the same operations in our simulations, we obtained estimated values of *κ* = 150−230kPa. This agreement is quite significant as we derived *κ*, a global mechanical property, by using parameters inferred from local indentation measurements.

**FIG. 3.**
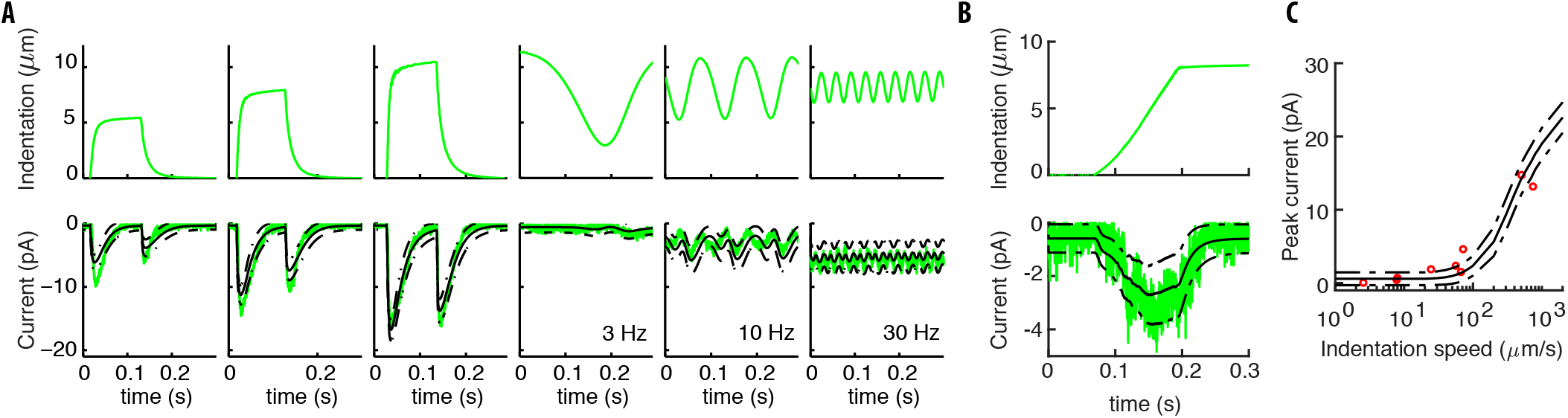
Our model captures experimental neural responses to various stimuli. (**A**) The applied experimental indentation (top); TRN’s response (bottom, green) and average predictions (solid black). Dot-dashed black lines correspond to one standard deviation above/below the mean. Experimental stimuli and neural responses are from Ref. [19]. (**B**) A typical ramp-like profile of indentation (top) and the corresponding current (TRN’s response in green; black lines as in panel A). (**C**) The predicted peak current *υs* the slope of the ramp for a total fixed indentation of 8*μ*m. Red circles indicate experimental data from Ref. [19].

### Experimental validation of neuron activation model

We now combine mechanics with the model for channel activation, extract predictions for TRN currents and compare to experiments. Predictions are valid for all profiles of stimulations, including new profiles not considered in [19], which highlight the importance of the full modeling developed here.

A detailed discussion of micro-cantilever systems of stimulation, and the *in vivo* patch clamp system to record neural responses was presented in [19]. A summary, together with specific differences for the data first reported here, is in Materials and Methods. Instances of stimuli and neural responses from [19] are shown in Fig. 3 and neural responses to new, pre-indented profiles are shown in Fig. 4.

Let us then describe how model predictions are obtained. Pressure *p* and the parameters of the mechanical part are as in the previous Section. Positions of the channels along the TRNs were randomly generated according to the log-normal experimental distribution [30], and results were averaged over those statistical realizations. As for the elastic filaments described by Eq. (5), their length is rescaled to unity as discussed in Appendix L, while their initial direction **Γ̂**(0) is distributed randomly. Namely, directions were generated uniformly in the semisphere with a non-negative component along the local outward normal **ŵ**_3_ to the neural membrane. Based on the deformation field determined by Eq. (2), we used Eq. (5) to compute the force on the channels as a function of time, and obtained the dynamics of the channels via Eq. (6). More details on the fits and the resulting values of the parameters are in Appendix L.

Our results in Fig. 3 manifestly capture the symmetric and rapidly adapting response of TRNs. The response to sinusoidal stimuli roughly oscillates at twice the input frequency due to the onset-offset symmetry of the touch response. At high frequencies, inertia in the switch between open and closed states of the channels contributes to the reduced amplitude of the oscillations. The response to ramps is intuitive: the slower the indentation, the smaller the response because of adaptation. Quantitative aspects are also well captured by our predictions.

Appendix M presents the histogram of the errors for individual realizations, which shows that neural responses are captured at that level as well (not just the mean, as in Fig. 3). The histogram also shows that restricting filaments to be initially or permanently tangential (**Γ**(0) · **ŵ**_3_ = 0 or x(*t*) · **ŵ**_3_ = 0), further improves results. The latter restrictions being speculative at this stage, we shall focus on the unrestricted model; we only note that tangential restrictions admit plausible molecular mechanisms, e.g. by microtubules that run along TRNs [40], which are attached to the neural membrane through filaments and impact touch sensation [41].

Additional insight is gained by delivering stimuli alternative to the classical profiles in Fig. 3, namely pre-indented stimuli in Fig. 4. Panel A contrasts standard and pre-indented steps, i.e. where an initial step (5*μ*m in our data) is delivered. The neural response to two steps of equal amplitude, one pre-indented and the other not, is substantially stronger for the former. That is surprising at first, since the amplitude of the steps is identical, and enough time between the successive half steps was left for adaptation. The explanation was obtained by using our model: it is indeed the case that channels adapt and return to their rest state; however, the tissue is deformed by pre-indentation, which leads to a more extended region of stimulation and more channels activated, as shown in Fig. 4E. The resulting predictions reproduce experimental trends, highlighting the importance of the coupling between mechanics and channel activation that constitutes the main focus of our paper.

### Explaining the effect of dissection protocols on neural responses

Experiments on *C. elegans* touch response are performed on dissected animals so as to access the TRN cell body in order to measure their currents. Two dissection procedures were employed in [19], which differ in having parts of the gonad and the intestines either removed (soft) or not (stiff). Names stem from the force-indentation curves in Fig. 5A, which evidences that the latter procedure better preserves the body’s integrity. Most experimental recordings reported here are for “soft” worms while data for “stiff” worms in Fig. 5A were used to predict the ratio *p*/*E* (Fig. 2C). Ref. [19] empirically showed that neural responses for soft and stiff worms are similar for displacement-clamped stimulations, while they strongly differ for force-clamped protocols. This Section analyzes the mechanical consequences of the above procedures, their effects on neural responses, and explains the empirical observation in Ref. [19].

Since internal organs of soft worms are removed away from the stimulation point, it is plausible that the dissection affects the internal pressure and has weaker effects on the indented external shell. This suggests to conservatively keep (in our model) the Young’s modulus *E* and the thickness *t* fixed, and modify *p*. Results of the corresponding simulations are shown in Fig. 5A. The slope of the force-indentation relation decreases with *p*; the best fit for soft worms is *p* = 1.6kPa, which is ∼ 4% of the value for stiff worms. Finally, the scatter around the mean shows that the more invasive dissection procedure results in stronger variability, with the corresponding *p* ranging from 0.04 to 16 kPa.

**FIG. 4.**
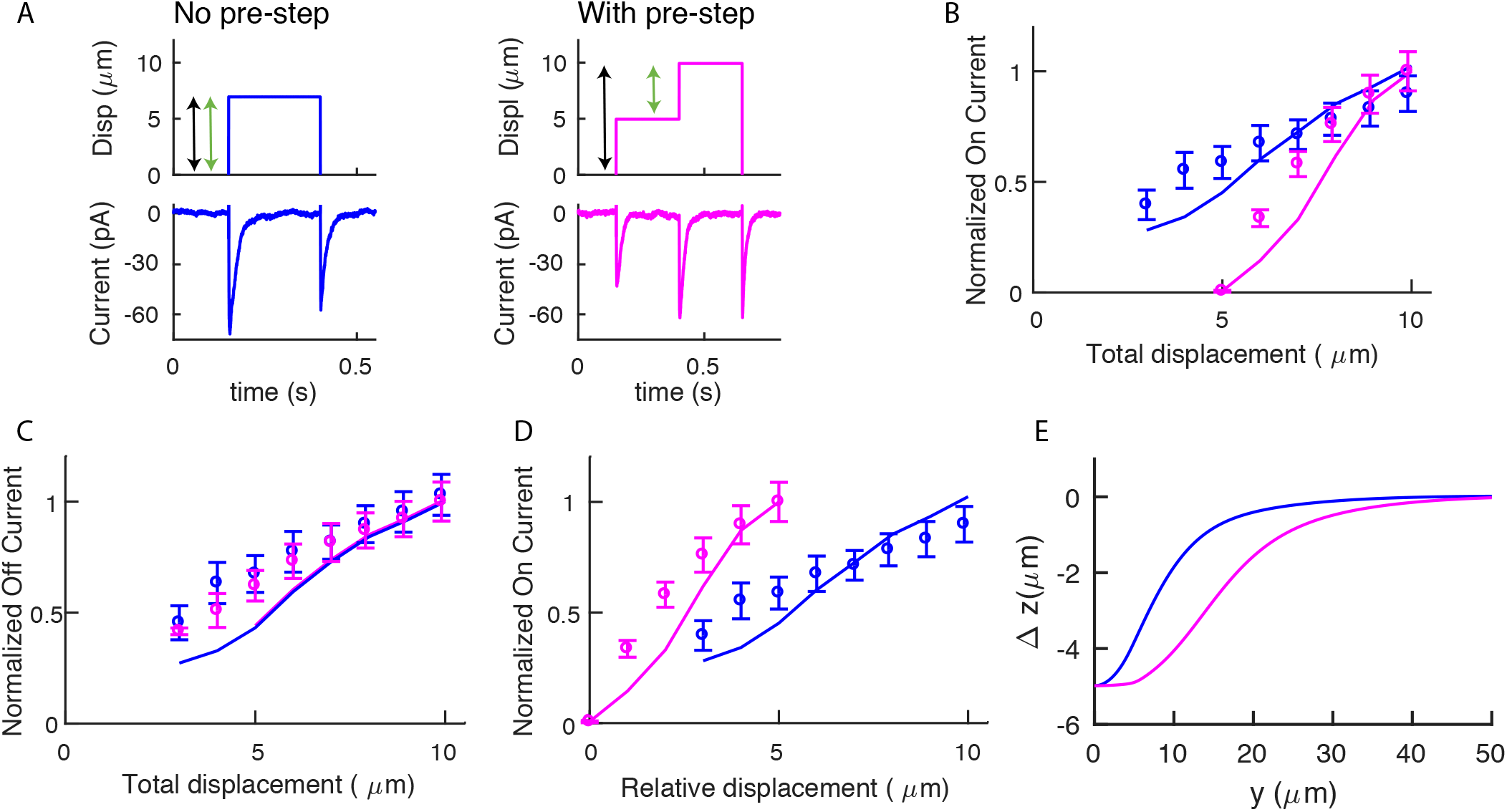
Pre-indented steps yield stronger responses due to their more extended deformation profile. (**A**) Stimuli delivered for standard (blue) and pre-indented (purple) steps. Black arrows indicate the total displacements for the on-currents in the following panels (colors match). Green arrows indicate relative displacements. Experimental stimuli and neural responses from ALM neurons were obtained as detailed in Materials and Methods. We recorded from 11 separate worms with 3 - 11 presentations of each stimulus per recording. Recordings were only included if they met the criteria outlined in the Data Analysis section of the experimental methods, which led to a final number number of biological replicates per displacement point that varied from 5 to 11. Representative traces shown here are from one biological replicate. (**B**) The on-current *υs* the total displacement (the pre-indentation for the purple points is 5*μ*m). (**C**) Off-currents are statistically indistinguishable, as expected since off-jumps are identical and adaptation erased the memory of the pre-step. (**D**) The on-current *υs* the relative displacements. Note the stronger response for pre-indented stimuli. (**E**) Changes in the profile of deformation: Δ*z* is the difference between the deformations after and before the (relative) stimuli. Note the greater extension for the pre-indented case, which is the reason underlying results in panel **D**. Open circles in panels B-D were normalized to the maximal currents detected and show the mean ± s.e.m.

To further analyze the effects of the dissection procedure, Fig. 5B shows the longitudinal profiles of vertical deformation for soft and stiff worms. The point is that the curves differ much less when displacement is clamped, rather than force. That is translated into predictions for currents as follows. We assume that the dissection procedure does not affect the channels and calculate their respective currents as described previously. Specifically, we fix the distribution of the channels, change *p* to the value corresponding to soft or stiff worms, and compute the neural response to force or displacement-clamped stimuli. Results are shown in Fig. 5C: responses for force-clamped stimuli widely differ for soft and stiff animals, yet they are similar for displacement-clamped stimuli. In sum, empirical observations reported in [19] are explained by the mechanics of the nematode and its coupling to neural activation.

Finally, we address the variability in Fig. 5C among soft worms, which we tentatively related to *p* varying over three orders of magnitude. For further support, we tested whether the observed variability could indeed be reproduced by keeping all parameters fixed but p. Results in Fig. 5D show that the peak current increases systematically with *p* for displacement-clamp and has the opposite behavior for force-clamped stimuli. The predicted change is larger in the former case, which is consistent with differences between soft and stiff worms. The model predictions are compared with an experimental dataset of 21 worms obtained in [19]. For each worm we inferred *p* from the force-indentation relation, pooling together animals with similar trends. Fig. 5D supports the initial hypothesis that changes in *p* among dissected animals are a major component in the observed variability.

**FIG. 5.**
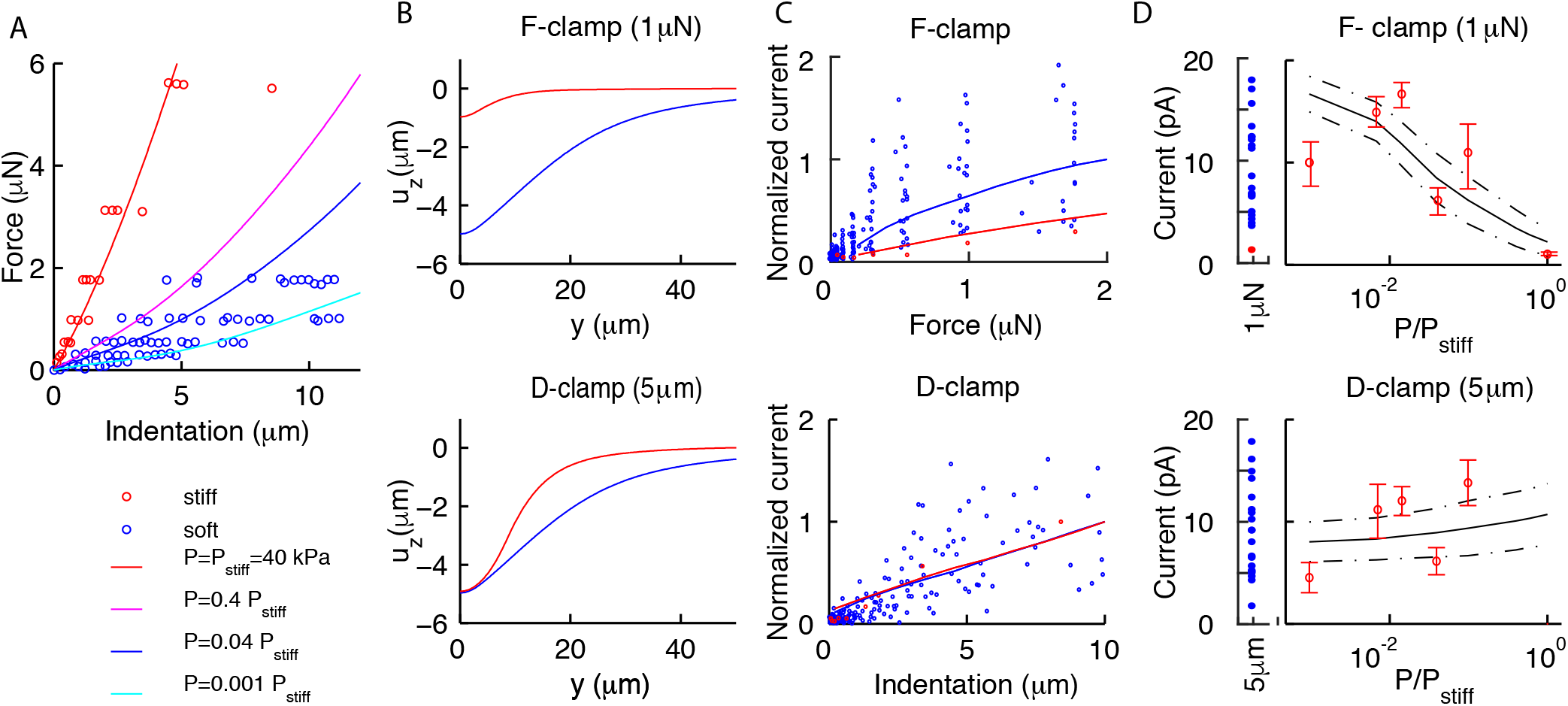
Our model explains previous experimental observations in Ref. [19] on differences among soft and stiff worms. (**A**) Experimental data (dots) and the average theoretical prediction (lines) for force-indentation relations. Best fit of pooled data for soft worms gives *p* = 1.6kPa; individual values are variable, with estimated *p* in the range 0.04 − 16kPa. (**B**) The vertical deformation profiles *u_z_ υs* the position *y* along the longitudinal axis for stiff (red) and soft (blue) animals. Note the widely differing profiles for the force-clamped curves. (**C**) Experimental (dots) and theoretical (mean value as continuous lines) peak current for force (top) and displacement-clamped (bottom) stimuli. The current is normalized by the mean peak in soft and stiff worms, respectively. (**D**) Peak current *υs* the pressure *p*, which shows that the model (continuous lines are the mean; dot-dashed lines are above/below one standard deviation) captures experimental trends (dots). Experimental data reproduced from Ref. [19] and derived from 4 and 21 recordings in the stiff (red) and soft (blue) conditions, respectively.

## RESULTS (ADDITIONAL MODEL PREDICTIONS)

In addition to the above predictions that could be tested against existing experimental data, our model yields additional results, which we present hereafter.

### Shell bending is weak compared to stretching; stiffness is dominated by internal pressure

The mechanics of pressurized shells relies on the balance between the internal pressure *p*, bending and stretching of the shell. Contrasting results have left undecided the previous balance for *C. elegans* [24, 37]. Here, we exploit our model to clarify this issue.

We computed the vertical deformation for different values of *p*/*E* and *t*, as previously done for the validation of the mechanical model. The longitudinal extension is quantified by the distance *y_h_* for the deformation to reduce to half of its maximum value (at the center of the bead). Fig. 6 shows that *y_h_* decreases, *i.e*. the deformation is more localized, when *p*/*E* increases. Conversely, as *t* reduces, the deformation is wider if *p*/*E* ≲ 10^−4^ and narrower if *p*/*E* ≳ 10^−4^.

To gain insight regarding the consequences of Fig. 6, we can use the thin-shell limit of Eqs. (2) in Appendix D. Reducing the limit equation to non-dimensional form as previously done for spheres [29], we find that the bending term is multiplied by the factor 1/*τ*^2^ ≡ *E*^2^*t*^4^/*p*^2^*R*^4^. If τ ≫ 1, the bending term is small, and (with the possible exception of boundary layer regions) the only remaining dependence on *t* is via the stiffness *S* ≡ *Et*. These arguments suggest to plot *y_h_ υs p*/*Et* as in Fig. 6: curves with different *t* indeed collapse for the values of *p*/*Et* that are relevant for experiments. We conclude that internal pressure and stretching of the shell provide the dominant balance.

We next compared the elastic energy of the shell with the work by the external forces. Results in Fig. 6C show that their ratio reduces as *p*/*E* increases, and the elastic energy tends to become marginal, which illustrates the dominance of the internal pressure in the body stiffness.

**FIG. 6.**
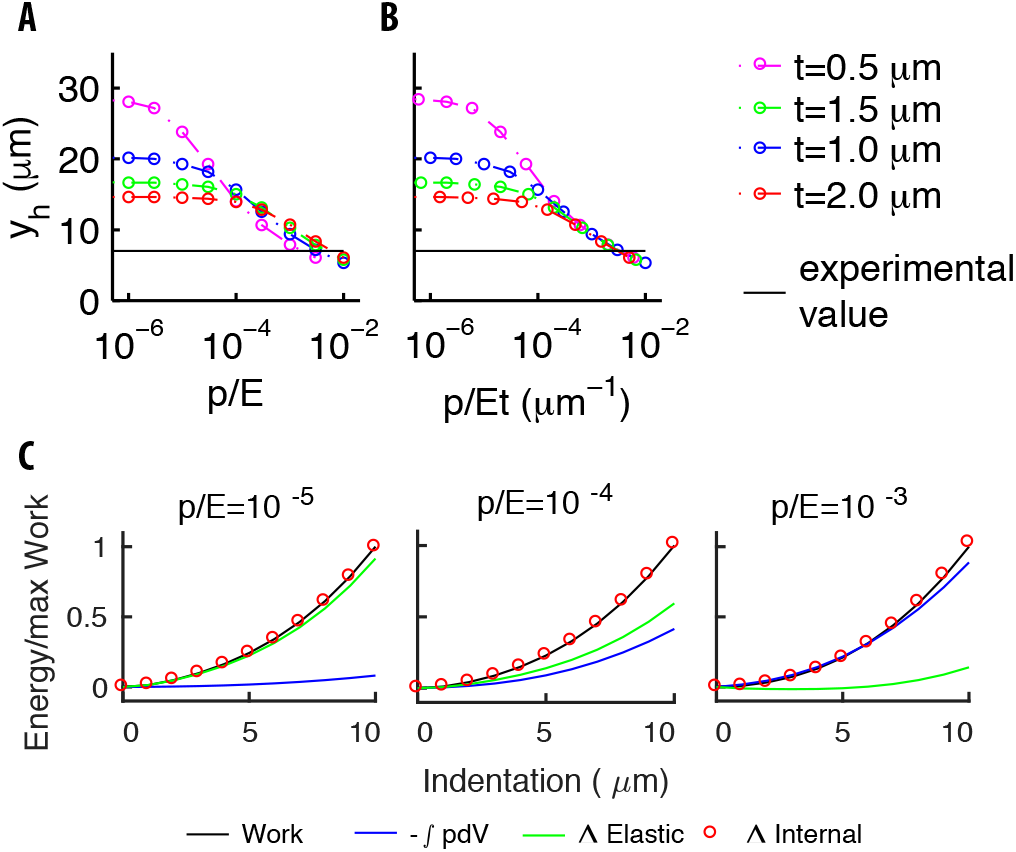
The mechanical balance for our model of pressurized shell. (**A**) The longitudinal extension *y_h_ υs p*/*E* for various thicknesses *t*. Different trends at small and large values of *p*/*E* reflect the contributions of bending to the elastic energy. (**B**) *y_h_ υs p*/*Et*. The collapse of the curves at the right end reflects the small value of the bending term coefficient (see the text). The value of *y_h_* found experimentally (black line) is well inside that asymptotic region. (**C**) The various contributions to energy, and the work done by the indenter for increasing values of the ratio *p*/*E*.

### Dependence of mechanical and neural responses on the radius of the indenting bead

**FIG. 7.**
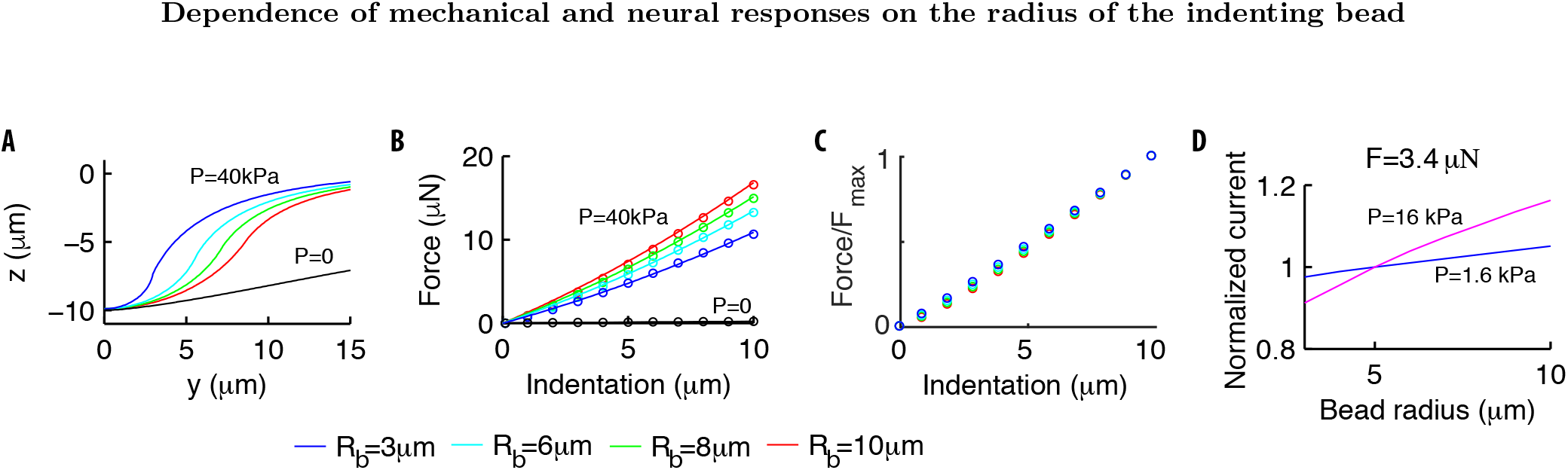
Effects of the radius Rb of the indenting bead. (**A**) In pressurized shells, the deformation profile depends on *R_b_* (colored lines) while the dependence disappears in the absence of internal pressure (black line). (**B**) The force-indentation relation for *p* = 40*kPa* and various *R_b_*. In strongly pressurized shells, the relation (colored dots) follows Eq. (12) (solid lines). For shells with *p* = 0, the curves collapse onto a unique curve (black dots). (**C**) The ratio *F* (*w*_0_)/*F_max_* of the force normalized by its maximum value is essentially independent of *R_b_*. (**D**) The peak current increases with *R_b_* in a *p*-dependent manner (the same holds for the sensitivity to the stimulus). The current is normalized by its value for *R_b_* = 5*μm*.

Experiments show that neural currents depend on the radius *R_b_* of the indenting bead [11]. Our aim is to quantify and predict this dependence from our model.

Results are shown in Fig. 7: *R_b_* influences both the deformation profile (Fig. 7A) and the force-indentation curve (Fig. 7B) for *p*/*E* in the experimentally relevant range. The curves are intuited as follows. The curvature of the deformation field at the indentation point increases with p/E until it matches the radius *R_b_*. As *p*/*E* increases further, the shell cannot become any steeper (the bead is rigid), so that it adapts to the bead in the contact region (see Fig. 2B), which widens with *R_b_*. The radius *R_b_* also controls the deformation outside of the contact region, namely the mid-maximum extension of the deformation (∝ R_b_, data not shown). As *p*/*E* reduces, the deformation becomes shallower at the indentation point and the role of *R_b_* vanishes (see Fig. 2B).

Similarly, Fig. 7A shows that the volume of the body to be deformed increases with *R_b_*, and more work is needed for a given maximum indentation *w*_0_. In formulæ: the work *Fdw*_0_ by the indenter roughly balances the contribution of the internal pressure −*pdV* (the elastic energy is small, as discussed previously) and the force-indentation relation is then given by *F* ∝ −*pdV*/*dw*_0_.

Qualitatively, larger *dV*’s associated with larger beads yield the nonlinear dependence in Fig. 7B. Quantitatively, we write *F* = *p*χ, where χ = −*dV*/*dw*_0_ has the dimension of a length squared and depends on *w*_0_, the shell and bead radii *R*, *R_b_*, and the ratio *p*/*E*. Keeping the latter fixed, we investigated numerically the behavior of χ and observed that the ratio *F*(*w*_0_)/*F_max_* does not depend on *R_b_* (see Fig. 7C). It follows that the dependence on *R_b_* should factorize out: 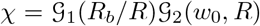 where 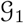 depends on *R_b_*/R for dimensional reasons. The function 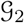 brings the length squared dimensionality and, in the limit of small *R_b_* and large p, behaves as *R w*_0_ [42]. It follows that 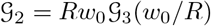. The functions 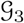 and 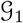 are empirically extracted from simulations as:

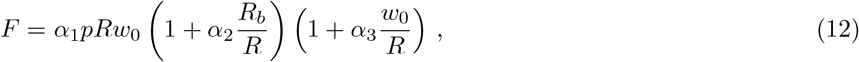

with α1 = 0.76, α2 = 2.1 and α3 = 0.66 for *p*/*E* = 0.01. Variations between *R_b_* = 3*μ*m and *R_b_* = 10*μ*m are on the order of few *μ*N, hence they should be accessible experimentally. Eq. (12) generalizes the linear relationship, valid in the limit of very small *R_b_*, obtained in [42].

Consequences for neural responses are in Fig. 7D. In agreement with [11], the peak current increases by ∼ 20% as *R_b_* goes from 3 to 10*μ*m, hence our prediction could be tested experimentally. A quantitative comparison with data in [11] was hampered by the lack of force-indentation measurements in [11], preventing us from inferring *p*.

Finally, it is worth remarking that the bead size also affects the dependence of the response on the circumferential position of the TRN. Indeed, Fig. 1 in Appendix F evidences that the profile of deformation decays rapidly as one moves circumferentially from the north pole (where the bead is indenting the body) toward the equator. That implies an appreciable dependence on the angular position of the TRN with respect to the bead, which will be stronger for smaller beads as the extension of their deformation is reduced.

### The onset/offset symmetry

Responses to the application/release of a step stimulus are comparable and well captured by our model. The goal of this Section is to further analyze the origin of this symmetry, by calculating the stimuli upon the channels and analyzing differences among microscopic gating mechanisms that are consistent with the symmetry.

A first key remark, which generally applies to thin shells [20, 22], is that the off-diagonal components *ε_xz_* and *ε_yz_* are small. Indeed, those two terms are proportional to the corresponding components of *σ*, which vanish due to the thinness of the shell (see [20, 22]). It follows from the definition of *ε* (see Appendix H) that vectors initially tangential and perpendicular to the surface of the cylinder, remain orthogonal even after deformation.

In addition to the general above property, the component *∊_xy_* is also negligible when the indenting bead is applied on top of the cylinder. The strain tensor is then diagonal, as confirmed by Fig. 8B.

The force acting on a single channel, as defined by (9) and calculated using (5), is shown in Fig. 8C. The force is maximal if the elastic filament is initially in the tangential plane while orthogonal filaments generate negligible forces. Notably, forces for tangential filaments have opposite signs yet very similar amplitudes at the onset and offset. The relation between vertical and tangential directions is key to the onset-offset symmetry and stems from the above discussion on thin shells. That constitutes the physical reason for our positing that tangential stimuli gate the channels: the orthogonal dynamics is indeed affected by the neural membrane, which *a priori* prevents any symmetry between inward and outward extensions.

### Neural response at the single channel level: isotropic vs directional models

Profiles in Fig. 8 are poised to discriminate between different microscopic models. Namely, alternatively to the isotropic choice in Eq. (8), we could consider the “directional” model with the preferential direction **v**:

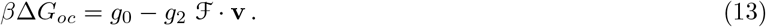

Contrary to Eq. (8), Eq. (13) breaks the symmetry for individual channels (see Fig. 8D), which can be restored though for the total current if channels along the TRN have their directions **v** in (13) independently and isotropically distributed (Fig. 9A).

A more quantitative analysis leans toward the symmetric model. Indeed, for the experimental value [11, 32] of the single-channel current *i*_0_ = −1.6±0.2pA, Eq. (13) underestimates the mean current (Fig. 9A). More generally, we can optimize parameters and calculate the errors in the fits of the experimental datasets: the probability distribution for Eq. (13) is broader and shifted to higher errors with respect to the isotropic model (see Fig. 9B). Similar conclusions hold if i0 is allowed to vary (Fig. 9C).

**FIG. 8.**
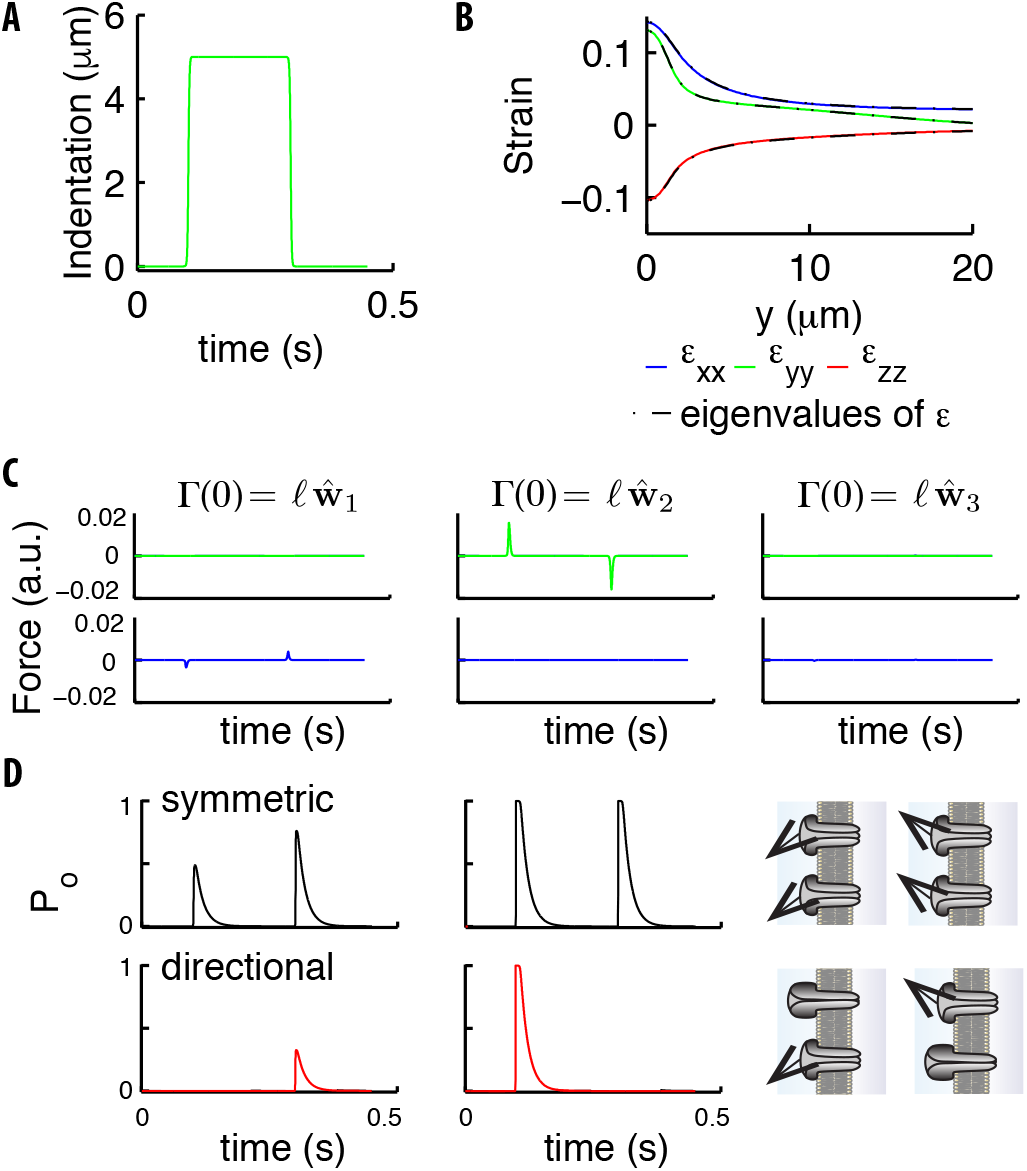
Stimuli on the channels due to a step. (**A**) Indentation profile for a step stimulus. (**B**) The diagonal components of the strain tensor *ε_ij_ υs* the longitudinal position along the cylinder. The overlap of those components (color) and the eigenvalues of *ε_ij_* (black) show that the tensor is essentially diagonal, which leads to the conservation of angles under deformation discussed in the main text. (**C**) The two components (green and blue curves) tangential to the neural membrane of the force acting upon on a channel (computed using Eqs. (5), (9)) for the stimulus in panel **A**. The panels refer to the different directions of the elastic filament (the first two are tangential and the third orthogonal to the neural membrane). (**D**) Gating probability for an individual symmetric or directional channel, as produced by the two tangential extensions in panel **C**. Parameters are: *y* = 1*μ*m, *θ* = 0, *υ* = cos(π/3) **ŵ**_1_ – sin(π/3) **ŵ**_2_. The sketch on the right illustrates that directional channels respond only to stimuli properly aligned with respect to their preferential direction while symmetric channels respond isotropically.

In sum, the analysis of available experimental data favors symmetric channels, but is not fully conclusive. New data will be needed, which is our motivation to describe hereafter two possible experiments.

A first approach relies on the noise level of currents. The intuition is that anisotropy reduces (for a given density of channels) the number of active channels along the TRN, and thereby leads to more noise. Specifically, the number of active channels could be inferred (Appendix J includes a generalization of the noise analysis in [11] to non-equally-stimulated channels) and compared to the number of channels measured by fluorescent tags [30]. Fig. 9D presents the Coefficient of Variation (CV) of the TRN current *υs* the stimulus strength, calculated over repetitions of a given stimulus. Differences in CVs are poised to permit discrimination and the approach described in Appendix J estimates that ∼ 100 trials suffice for their reliable measurement.

A second alternative exploits the architecture of *C. elegans* neurons: TRNs extend longitudinally for about half of the nematode’s length, leaving a region around its center that is relatively insensitive to touch (see [43]). Fig. 9E indicates the range over which effects of indentation are felt by individual channels; panel F shows the differences between microscopic models as the indenting bead slides along the longitudinal direction. An additional relevant statistic is the asymmetry between on- and off-currents. The logic is that, as the number of stimulated channels decreases, asymmetries should become more substantial if the channels are not isotropic, see Fig. 9G. An appealing possibility is the stimulation of the worm in its center (negative coordinates in panels F,G). There, the number of activated channels is small, which could indeed bring microscopic insight.

**FIG. 9.**
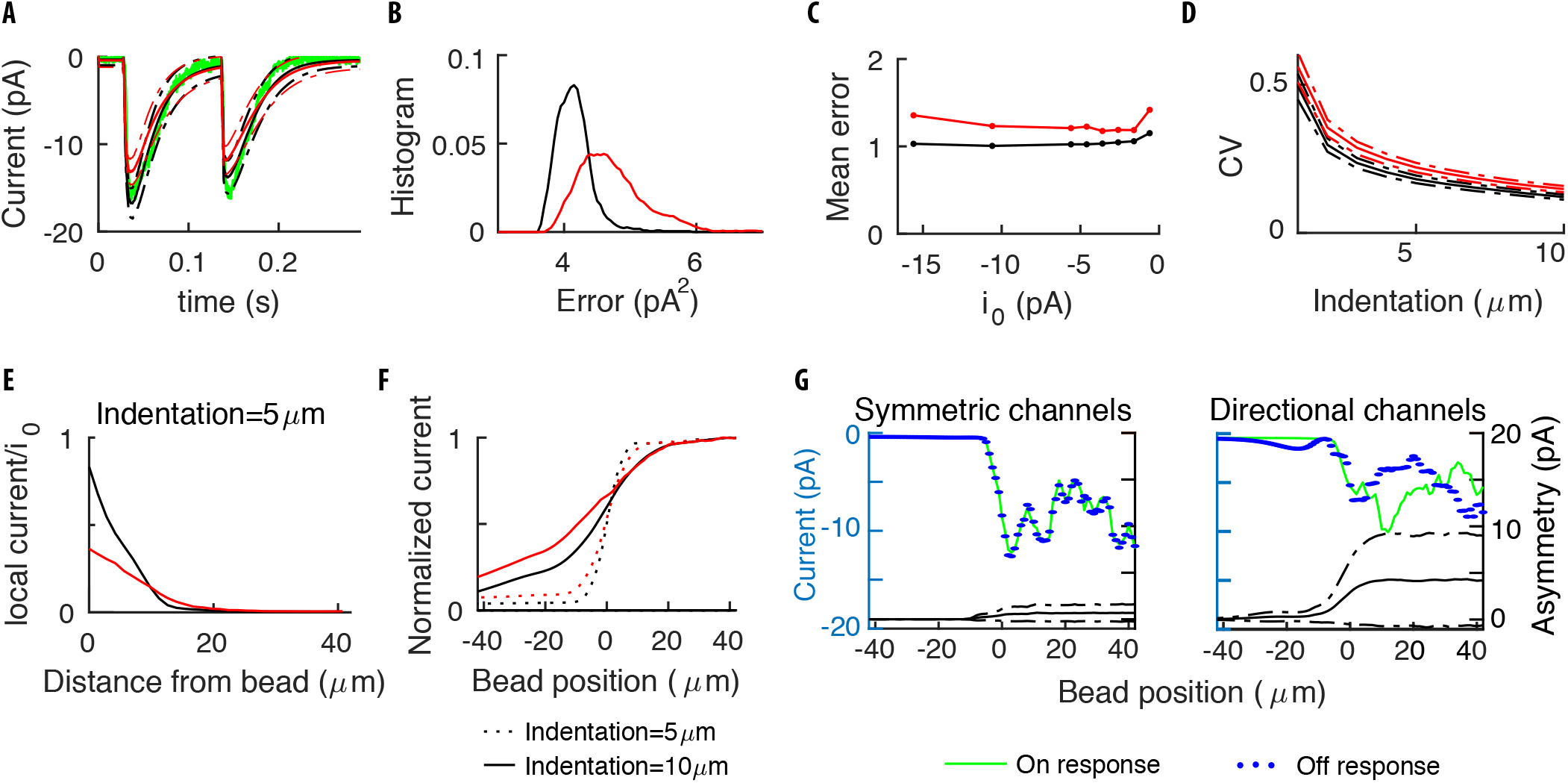
Symmetry of the single channel response. (**A**) Mean neural current response to a step, for symmetric (black) *υs* directional (red) channels. (**B**) Histogram of the errors in reproducing the data of Fig. 3 obtained with the two above models for different realizations of the channels’ distribution. Symmetric channels (black) give a better description. (**C**) The mean error as the maximum current *i*_0_ per channel is varied. (**D**) The Coefficient of Variation (CV) of the TRN current *υs* the stimulus strength, calculated over many repetitions of a given stimulus. (**E**) The average current for a channel (normalized by its maximum value *i*_0_) as a function of its distance to the center of the indenting bead. (**F**) The current flowing along the TRN *υs* the position of the indenting bead. The origin indicates the end of the TRN; negative coordinates correspond to the relatively insensitive zone in the middle of the body of the worm. (**G**) The colored curves show the predicted current for symmetric and directional channels, for a given distribution of the channels. The black curves show the expected level of asymmetry between onset and offset, as quantified by the standard deviation 〈(***I**_on_* – ***I**_off_*)^2^〉^1/2^ between the peak responses ***I**_on,off_* at the onset/offset of the stimulus, averaged over the distributions of the channels. Dashed-dotted curves show the range of expected asymmetries in individual realizations.

## DISCUSSION

We presented the first comprehensive description of *C. elegans* mechanics and activation of TRNs. That allowed us to compute the TRN currents, which result from the response of mechanosensitive channels to stimuli applied on the skin of the worm. Predictions were compared to in vivo patch-clamp neurophysiology experiments.

Our model explains several aspects of the coupling between mechanics and neural responses. First, we capture the experimental current flowing along TRNs in response to a wide range of indentation profiles, including new preindented ones. Second, the model explains the empirical observation previously made in Ref. [19] that neural responses for soft and stiff worms are similar for displacement-clamped stimulations, while they strongly differ for force-clamped protocols. The cause is traced back to the mechanics of the skin deformation. Third, the model rationalizes how the response is modified by the dissection procedures and shows that internal pressure of the nematodes is a major source of variability among different animals. Fourth, the model predicts how the neural response should change with the size of the indenter. Finally, the variation of the deformation with respect to the circumferential variable evidences an appreciable dependence on the angular position of the TRN with respect to the indentation bead. It would then be important to design the set-up of future experiments so as to make that information available.

A by-product of our analysis is that it provides insight on two points related to the mechanics of the worm.

First, our results reconciled conflicting results [24, 37] on mechanical properties in wild type and mutants. Specifically, we showed that the mechanical response of the nematode is captured by an elastic cylindrical thin shell in a pressure-dominated regime. The theory makes testable predictions on the dependence of the force-indentation relation on the indenter size, as well as the effects of mutations in the cuticle on the bulk modulus. An experimental verification of those predictions would further support the major role of internal pressure in *C. elegans* body mechanics.

Second, the fact that our model yields best results for boundary conditions with no force at the two ends of the cylinder, suggests that the body of the nematode might relax longitudinal components of the stress. One possible mechanism is through the annular structure of the cuticle [23], which may be effectively described as a shell with anisotropic Young’s moduli. The relaxation of longitudinal stresses could be relevant for the motility of the nematode since it facilitates its bending, an ingredient necessary for swimming [44, 45].

We have shown how neural responses have similar amplitude when the stimulus is applied or released, which addresses the long-standing puzzle of the onset-offset symmetry. Our gating mechanism highlights the importance of components of the stimulus tangential to the neural membrane. That fundamentally differs from previous models [1, 14, 15], which involve orthogonal components. At the microscopic level, we inquired whether the symmetry holds for individual channels or at the population level only (individual channels are asymmetric yet their preferential directions are randomly distributed and their cumulative effect is again symmetric, as suggested for *Drosophila* sound receiver [46]). We showed, however, that a model with symmetric channels gives a better description of existing data, hinting at symmetry for single channels. Definitive evidence could be obtained by the experiments that we suggested in Fig. 9C and G, with the noise levels better controlled and the stimulation point moved along the longitudinal axis so as to assay a variable number of channels. The ideal experiment would be to precisely assay the neural response to stimuli in the central dead-zone of the body, where few channels are likely to be directly stimulated (see [43]).

In our current description, we assumed that the material composing the shell is purely elastic and the dependencies on frequency result from the gating of the channels. While this procedure successfully captures many experimental observations, it is known that tissues do feature viscous effects [36]. Future developments will address viscoelastic effects, which should be relevant to the understanding of touch sensation at high frequencies.

Finally, it is worth noting that our modeling ultimately relies on the fact that touch receptor neurons are close to the surface of the skin’s thin layers. This leads to physical effects peculiar to thin shells, namely the importance of tangential forces, which are at the basis of the gating mechanism discussed here. Since the above features are common in touch sensation, we expect results and methods that we developed to be widely relevant.

## MATERIALS AND METHODS

We incorporated most of our modeling methods in Results and Discussion. For additional details, please see Appendices. Experimental methods are found hereafter.

### Experimental methods

#### Nematode strains

The following transgenic *C. elegans* nematodes were used: TU2769 *uIs31[mec-17p∷gfp]* III [11] and TP12 *kaIs12[col-19∷gfp]* [47]. The *uIs31* transgene expresses GFP exclusively in the TRNs, enabling in vivo recordings from these neurons and the *kaIs12* transgene encodes a fusion between the COL-19 collagen protein and GFP, labeling cuticular annuli. Animals were grown on OP50 at either 15°C (TU2969) or 20°C (TP12) and used as well-fed L4 larvae or young adults.

#### Imaging Cuticle Deformation

TP12 worms were immobilized with 0.1μm polystyrene beads on a 6% NGM agarose pad. 10μm glass beads (Duke Scientific) for indenting the worms were spread onto a coverslip, which was inverted to cover the agarose pad holding the worms. To image the worms, we used a high-magnification camera (Orca-R2, Hamamatsu) on an inverted microscope (Leica) with an EGFP filter set and a high-numerical aperture 63x oil immersion lens, to yield a shallow depth of field ≈ 0.1μm for optical sectioning. When glass beads were trapped between the cuticle of the animal and the coverslip, we were able to capture fluorescence images of COL-19∷GFP in the cuticle at >10 different focal planes. At each focal plane, we measured the radius of the bead and the radius of the cuticle deformation (by identifying where the cuticle was in focus). We then calculated the depth of the plane based on the radius of the bead at the focal plane. Experimental data shown are a combination of all focal planes for 2 adult animals.

#### Electrophysiology

Worms were immobilized on 2% agarose pads with WormGlu (GluStitch), dissected, and patch-clamped as described in [19]. Recordings were performed on the ALMR neuron due to geometric constraints of the stimulator system; ALMR is bilaterally symmetric to the previously used ALML neuron. The extracellular solution contained (in mM): NaCl (145), KCl (5), MgCl_2_ (5), CaCl_2_ (1), and Na-HEPES (10), adjusted to pH 7.2 with NaOH. Before use, 20mM D-glucose was added, bringing the osmolarity to ∼325mOsm. The intracellular solution contained (in mM): K-Gluconate (125), KCl (18), NaCl (4), MgCl_2_ (1), CaCl_2_ (0.6), K-HEPES (10), and K_2_EGTA (10), adjusted to pH to 7.2 with KOH. Before use, 1mM sulforhodamine 101 (Invitrogen) was added to help visualize successful recording of the neuron.

Membrane current and voltage were amplified and acquired with an EPC-10 USB amplifier and controlled through Patchmaster software (HEKA/Harvard Biosciences). The liquid junction potential between the extracellular and intracellular solutions was −14mV and was accounted for by the Patchmaster software. Data were sampled at 10kHz and filtered at 2.9kHz.

#### Mechanical stimulation

For mechanical stimulation during patch-clamp electrophysiology, previous studies used either open-loop systems with a piezoelectric bimorph [11] or stack [41, 48–50] with no measurement of actual displacement or a closed-loop system with a stimulus bead at the end of a piezoresistive cantilever for force detection, driven by a piezoelectric stack [19]. Here, we use an open-loop system adapted from the piezoelectric stack system with a photodiode motion detector described in [51]. This enables faster stimulation than the force-clamp system [19, 52] at the expense of control over and measurement of exact force and indentation. The photodiode detector allows for a readout of the time course of the displacement of the stimulator.

An open-loop piezoelectric stack actuator with 20*μ*m travel distance (PAS-005, ThorLabs) was attached with marine epoxy (Loctite) to a 0.5” diameter, 8-inch length tungsten rod, and mounted on a micromanipulator (MP-225, Sutter) at a 17° angle to allow the stimulator to fit beneath the microscope objective.

For detecting probe motion at the 0.5-10*μ*m scale, we adapted the system from [51] to use the SPOT-2D segmented photodiode (OSI Optoelectronics), and mounted it in an XY translator on top of a rotation stage (ST1XY-D, LCP02R, ThorLabs) to enable alignment of the photodiode gap perpendicular to the direction of probe motion. This was affixed above a secondary camera port on the microscope (Eclipse E600FN, Nikon) with no additional magnification.

To create a defined and reproducible contact surface for the stimulation probe, we adapted the bead gluing technique used previously for the force-clamp system [19, 52], but with an opaque bead that allowed for a clear signal from the photodiode motion detector. Borosilicate glass pipettes (Sutter, BF150-86-10) were pulled and polished to a tip diameter of 10-15*μ*m, and 20-23*μ*m diameter black polyethylene beads (BKPMS-1.2, Cospheric) were attached with UV-curable glue (Loctite 352, Henkel). Pipettes with attached beads were trimmed to a length of 1-2cm, placed in the pipette holder, and waxed in place with sealing wax (Bank of England wickless, Nostalgic Impressions). A high-resolution 3D-printed acrylic pipette holder (custom design) was attached with marine epoxy to a steel tip (PAA001, ThorLabs) mounted on the piezo stack.

After cell dissection, but before making a gigaseal for patch clamp, the front edge of the stimulator bead was moved into place and visually aligned under the 60X objective with the highest visible edge of the worm’s cuticle at a distance of 108±36*μ*m anterior to the ALM cell body.

#### Stimulus control and data acquisition

All systems described here were controlled through HEKA Patchmaster software with a 10kHz sampling frequency. The voltage output from the EPC-10 amplifier (HEKA) was adjusted based on the total range of the stack for a relationship of 0.418 V/*μ*m. This command signal was filtered at 2.5kHz on an 8-pole Bessel filter (LPF-8, Warner Instruments) and then amplified with a high-voltage, high-current Crawford amplifier [53] to achieve a signal between 0-75V which was sent to the stack. The stack was biased with a starting offset of 3-4*μ*m, and the largest displacement used was 3-4*μ*m less than the upper limit of the stack’s travel distance, ensuring that stack motion was linear. The analog signal from the photodiode circuit was digitized at a rate of 10kHz by the EPC-10 amplifier and Patchmaster software, for temporal alignment of the probe motion signal with the evoked current response.

#### Data analysis

Whole-cell capacitance and series resistance were measured as previously described [54]. Data analysis was performed with MATLAB from Mathworks (data import and analysis functions are available online at: http://github.com/sammykatta/Matlab-Patchmaster) and Igor Pro (Wavemetrics).

Only recordings with holding current < − 10pA at −60mV and series resistance < 210MΩ were included in the analysis. Since the voltage was not changed during the course of these experiments, we did not correct for voltage errors due to uncompensated series resistance.

## ACKNOWLEDGMENTS

We thank Z. Liao for technical assistance with C. elegans animals. Experimental research was supported by NIH grants (R01-NS-047715, R35-NS-105092 to MBG; R01-EB-006745 to BLP and MBG) and fellowships (F31-NS-093825 to SK).

## Appendix A Numerical evaluation of the amplitude of the deformation gradients

In the main text, we exploited the property that gradients of the deformation field **u** (which are non-dimensional quantities) are small compared to unity. This was used first to justify a Hookean relation between stress and strain, and then in the derivation of the friction force acting on the elastic filaments. The goal of this Appendix is to provide an empirical *a posteriori* validation of this assumption.

Fig. 1 shows the various gradients components for an indentation of 10*μ*m, which is the greatest value in experiments. To provide an upper bound, we focus on the region of maximum deflection, i.e. along the longitudinal direction at the top of the cylinder. Over a wide range of positions, both in soft and stiff worms, the gradients are indeed smaller than unity. The only exception is the component *du_z_*/*d_y_*, which approaches unity in a small region around *y* = 5*μ*m, which is where the bead detaches. However, it is sensible to neglect even this contribution for predictions of the neural current. Indeed, values in Fig. 1 are an upper bound, and the region is just a few *μ*m wide, hence only a small number of channels are possibly concerned (we remind that the average inter-channel distance is ∼ 2*μ*m).

## Appendix B How does the radius of the cylinder change with the internal pressure ?

In this Appendix we discuss modifications in the geometry of a cylindrical shell upon application of internal pressure. We first compute analytically the deformation field in the linear approximation, and then extend the derivation in the nonlinear regime with the assistance of numerical analysis.

The 3D elasticity equations (Eq. (2) in the main text) in the linear approximation and cylindrical coordinates read

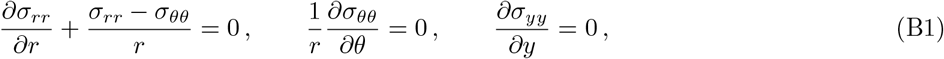

where *x* = *r cos*(*θ*), *z* = *r* sin(*θ*); non-diagonal terms of *σ* vanish due to the symmetry of the problem. Boundary conditions are

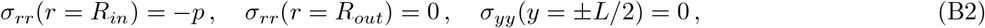

where *R_in_* = *R* − *t*/2 and *R_out_* = *R* + *t*/2. The condition of zero longitudinal stress at the two ends of the cylinder is motivated by the results in Appendix G.

**Appendix A, Figure 1.**
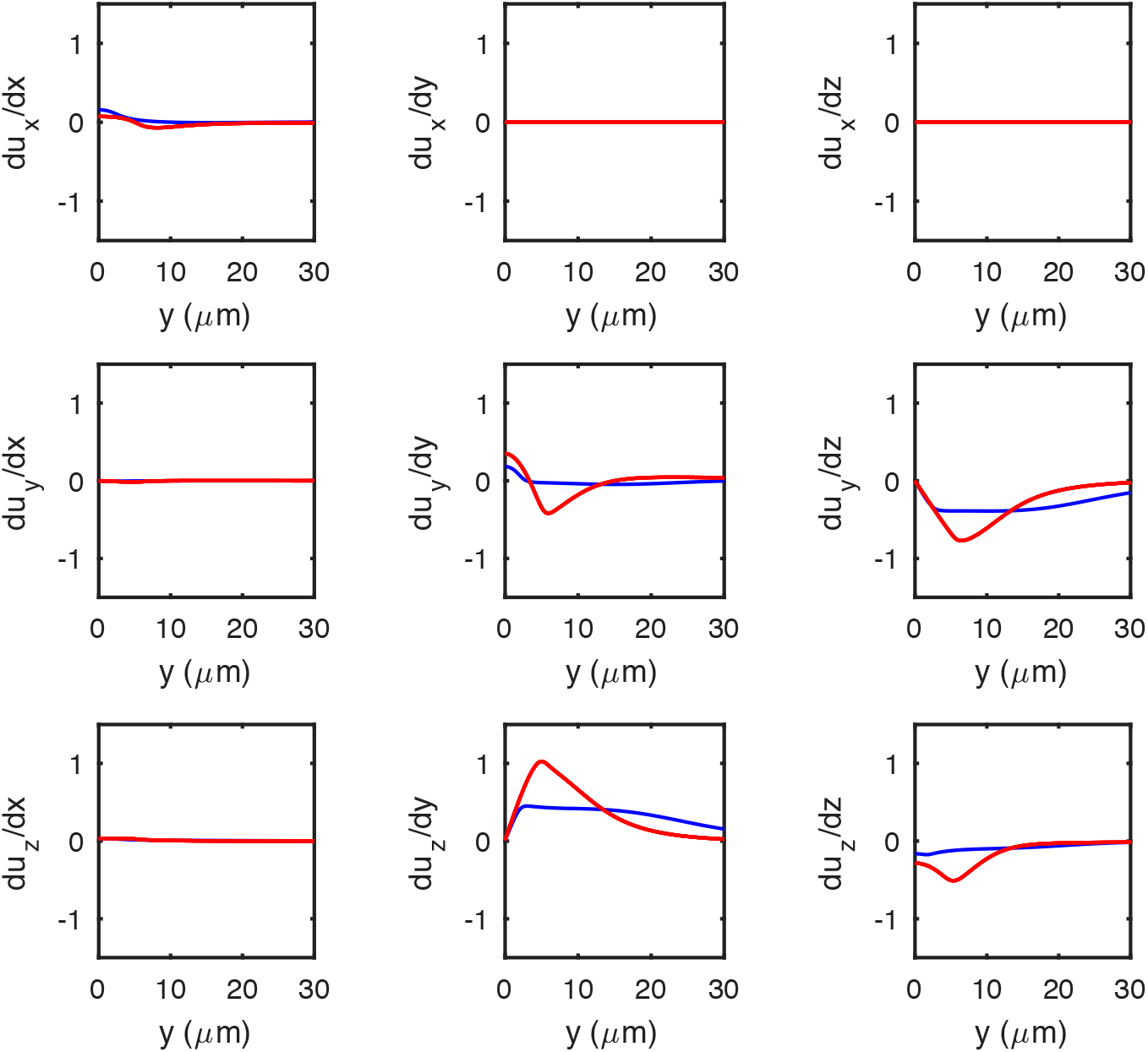
The gradients of the deformation along the longitudinal direction of an indented cylinder. Gradients of **u** computed by using numerical simulations for soft (*p*/*E* = 4 × 10^−4^, blue) and stiff (*p*/*E* = 0.01, red) shells. Note their moderate amplitude even for the indentation of 10*μm* shown here, which is the strongest that we consider.

The third line of Eq. (B1) and the boundary conditions imply *σ_yy_* = 0 along the shell. By using the constitutive Hookean relations between stress and strain tensor in the main text, we obtain

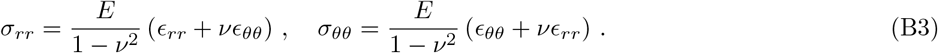

For small deformations, the diagonal components of the strain tensor (see Eq. (1) in the main text) are given by

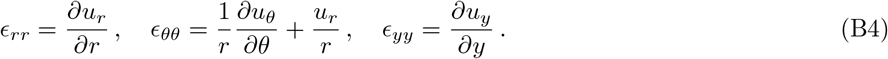

Due to the geometry of the problem *∂u_θ_*/*∂θ* = 0; by using Eqs. (B1), (B3) and (B4) we find the following equation for the radial deformation

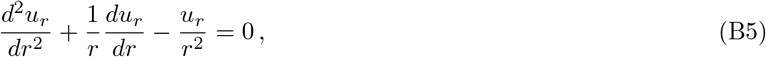

whose general solution is *u_r_* = *a*/*r* + *br*. Using the boundary conditions we obtain

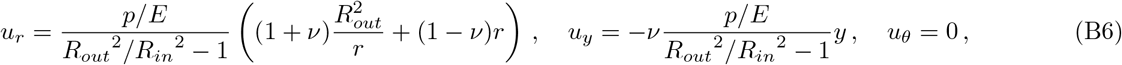

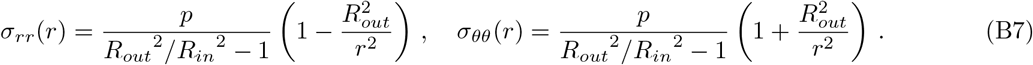

It follows from the solution (B6) that the relative change in radius, thickness and length, in the limit *t*/*R* ≪ 1, are given by

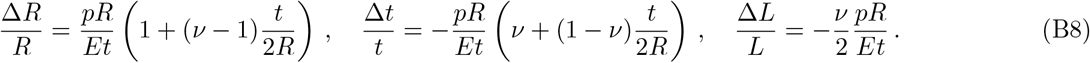

Fig. 1 demonstrates good agreement of these predictions with numerical simulations for small values of *p*/*E*. The corresponding expressions for the change in the volume *V* = π*L* (*R* + *t*/2)^2^ of the cylinder and its bulk modulus read

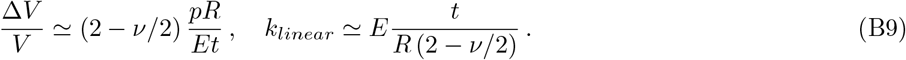

As *p*/*E* increases, nonlinear behaviors beyond the linear description of Eqs. (B1) and (B4) become important. An analytical description of the corresponding deformations would require to take into account nonlinear terms in the original Eqs. (1) and (2) of the main text.

For our purpose here, the following empirical approach will suffice. The linear solutions (B8) depend on two dimensionless small parameters: *pR*/*Et* and *t*/*R*. In fact, Eq. (B8) has *t*/*R* appearing only multiplied by *pR*/*Et*; that makes its contribution small, and implies that the functions depends on *pR*/*Et* only, at the dominant order. Fig. 1 shows that this property extends in the nonlinear regime: indeed, the curves for relevant values of *t*/*R* collapse when plotted *υs pR*/*Et*. Using this empirical observation, we looked for a power series in *pR*/*Et* and found numerically that the following functional forms describe the deformations in the regime relevant for our problem (*pR*/*Et* ≤ 0.4):

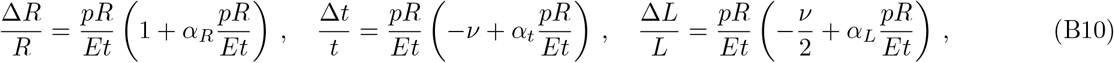

where *α_R_* = −0.6182, *α_t_* = −0.0626, *α_L_* = 0.0479. Finally, by using the relations (B10), we obtain the volume *V* and the bulk modulus *k* of the cylinder as a function of *p*/*E*, as shown in Fig. 1G,H.

**Appendix B, Figure 1.**
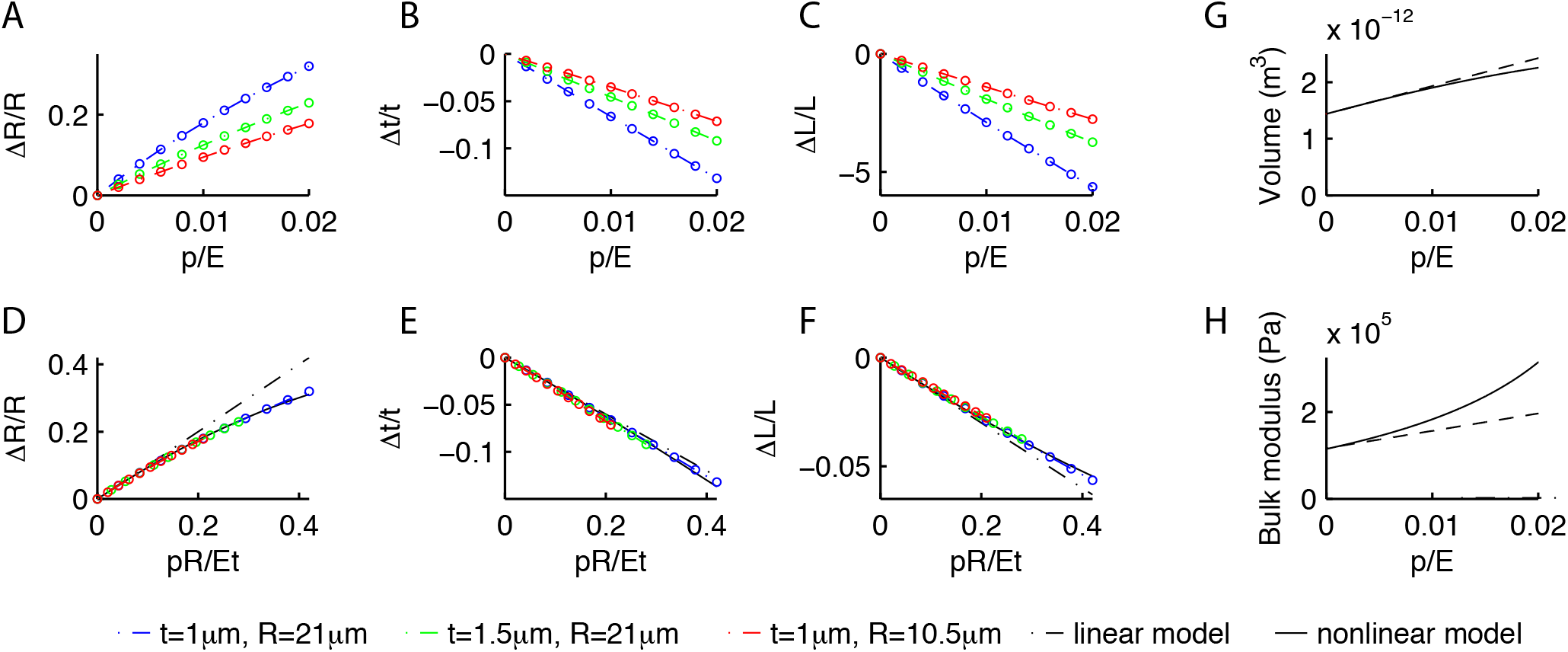
Deformations of a cylindrical shell due to internal pressure. Relative change in the radius *R* (A), thickness *t* (B) and length *L* (C) of the shell as a function of *p*/*E* and various *t/R*, as obtained from our numerical simulations. As *p* increases, *R* increases whilst *t* and *L* decrease. (**D,E,F**) While the curves in the previous panels change with *t/R*, the curves are collapsed by plotting them against *pR/Et*, which suggests that the contribution of terms in *t/R* is negligible. The collapsed behavior agrees with the linear prediction Eq. (B8) for *pR/Et* < 0.1 (dash-dotted lines), and is well captured by the empirical Eq. (B10) in the moderately nonlinear regime. Using the nonlinear Eq. (B10), we computed the change in volume (G) and bulk modulus (H) of the shell as a function of *p*/*E* (dashed lines are the linear predictions (B9) valid for small values of *p*/*E*).

## Appendix C Effects of external atmospheric pressure on mechanical and neural response

In the main text, we studied the mechanical properties of *C. elegans* body as a function of the pressure parameter *p*, i.e. the difference between the internal and the external atmospheric pressure *P_atm_*, neglecting the latter. This Appendix shows that our results hold also when *P_atm_* is taken into account.

To gain insight, we first adapted the calculation in Appendix B to the case where atmospheric pressure *P_atm_* is considered. The equations are not modified yet the boundary conditions on the internal and external surface of the shell become

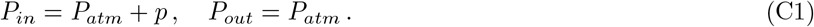

By following the same procedure detailed in the previous Section B, we obtain that the radial deformation of a pressurized shells is given by

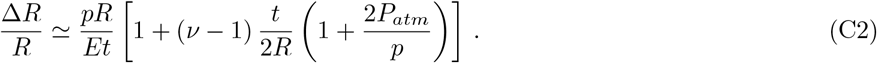

Eq. (C2) suggests that effects of atmospheric pressure on the mechanical response of the shell are negligible as long as *P_atm_t*/*pR* ≪ 1. To test this hypothesis, we simulated indentation experiments with and without atmospheric pressure for different values of p; results are shown in Fig. 1. As expected, for *p*/*E* ≈ 10^−2^, both the force indentation relation and the deformation profile are not modified by the atmospheric pressure. It follows that all mechanical properties and neural responses derived in the main text for stiff worms (where the internal pressure is high and *p*/*E* ≈ 10^−2^) are not modified by the inclusion of atmospheric pressure.

Soft worms were shown in the main text to have smaller values of *p*/*E* because of the dissection procedure. Fig. 1 shows that the force indentation relation does not change significantly yet the deformation profile becomes wider as *p*/*E* reduces. This modification is not relevant to describe mechanical properties of intact animals but it might *a priori* influence the neural response in soft worms. Therefore, we computed numerically the neural response to indentation experiments in soft worms with and without atmospheric pressure. As shown in Fig. 2, the performance of the model in describing neural data is not modified substantially, which shows that results of the main text are generally valid even when atmospheric pressure is taken into account.

**Appendix C, Figure 1.**
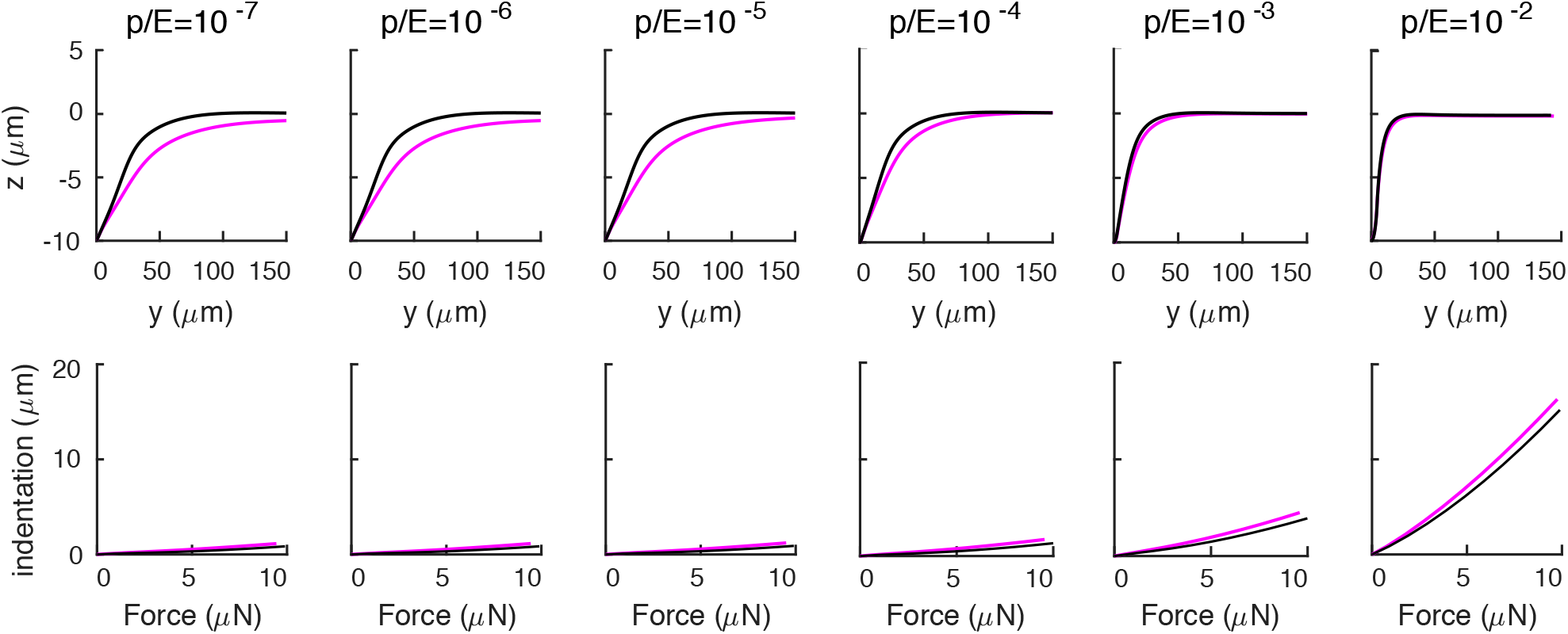
Effect of atmospheric pressure on mechanical response. Deformation profile (first row) and force indentation relation (second row) for a cylindrical shell (with the same properties as in the main text) with (purple) and without (black) atmospheric pressure.

## Appendix D The thin shell limit

In the limit when the shell is thin and the surface is shallow, the 3D elasticity Eqs. (2) reduce to [21, 22]:

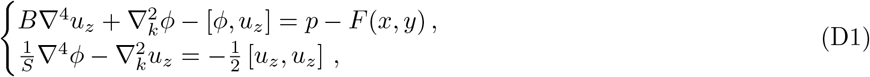

which is the limit equation that was used in the main text for analyzing the energetic balance between bending,stretching and internal pressure. The brackets in (D1) are defined as 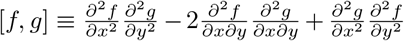, derivative 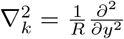. Eqs. (D1) are two-dimensional, with the variables *z* = *z*(*x,y*) and *u_z_* = *u_z_*(*x,y*) representing the middle surface and the deformation field of the cylindrical shell, respectively. The deformed surface is *z* + *u_z_* and we chose the axes so that the plane *z* = 0 is tangent to the top of the cylinder. The Airy stress function *φ* is the scalar function that parametrizes the in-plane components of the stress tensor as 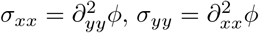 and 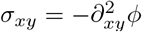.

**Appendix C, Figure 2.**
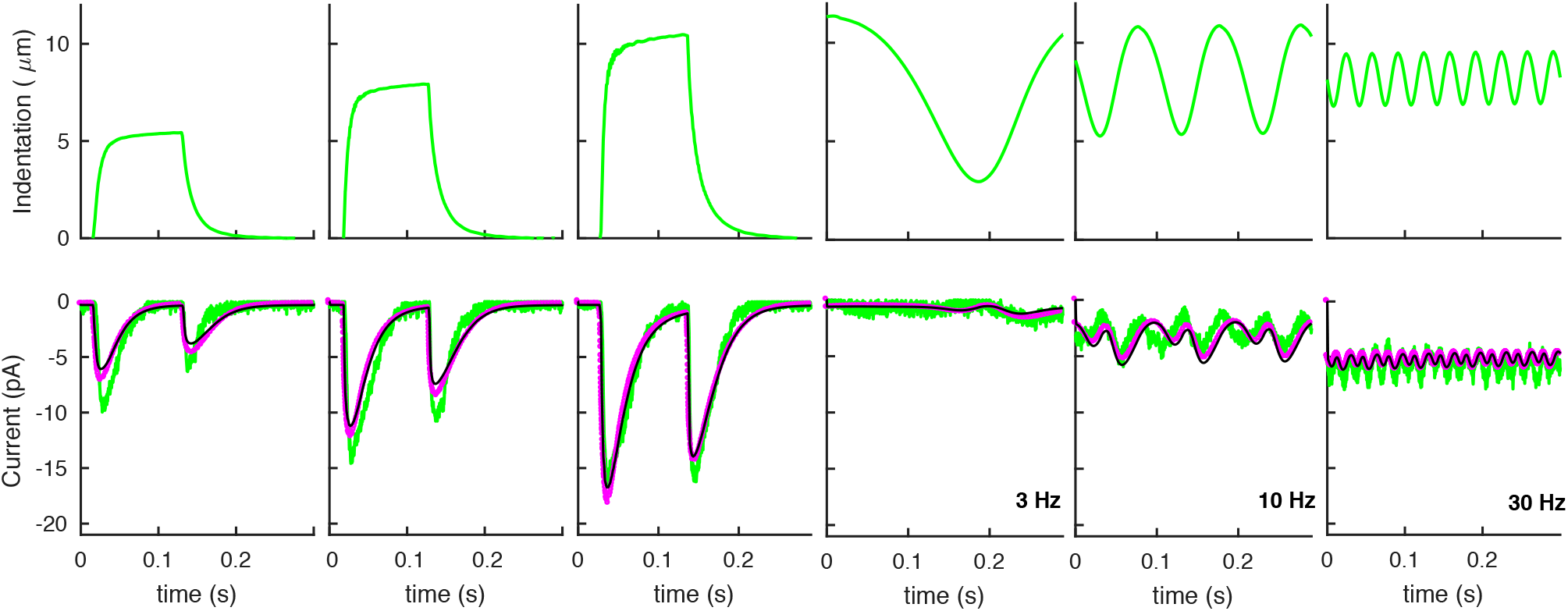
Effect of atmospheric pressure on neural response. Neural responses of the model presented in the main text to the experimental stimuli in Fig. 3 of the main text. Green curves are experimental data (as in Fig. 3 of the main text); purple and black curves are model predictions with and without atmospheric pressure, respectively. The ratio *p*/*E* is 7‰ of the value for stiff worms.

The above simplifications are due to the thinness of the shell and the resulting small vertical components of *σ*.

The parameters *B* = *Et*^3^/12(1 − *ν*^2^) and *S* = *Et* are the bending and stretching stiffness, respectively. Finally, *p* and *F* are the internal pressure and the external force applied by the indenter. In the limit *R* → ⋡, the 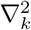 term is negligible, and Eqs. (D1) reduce to the Föppl-von Karman equations for a thin plate [21, 22].

## Appendix E Validation of the numerical scheme used to determine the mechanical response

In the main text, we described the mechanics of the nematode as an elastic cylindrical shell under pressure. Because of the geometrical nonlinearities involved, numerical simulations are the main tool available for determining the resulting mechanical response. The goal of this Appendix it to give more details on the tests that we employed to validate the numerical procedure discussed in the main text. Tests rely on elasticity problems that were previously investigated, namely small indentations of cylindrical shells (where an analytical solution is available [28]), and large indentations of pressurized spherical shells (where a simplified framework was derived in [29]). In all cases, agreement was obtained.

### 1. Small indentations of cylindrical shells

A thin cylindrical shell subject to equal and opposite concentrated radial loads was investigated in [28]. For indentations *w_0_* ≪ *t*, where *t* is the shell thickness, the equations of 3D elasticity (Eqs. (2) in the main text) reduce to [55]:

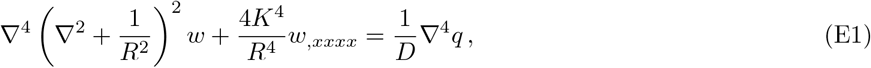

where *q* is the applied load, *x* and *θ* are the cylindrical longitudinal and angular coordinates, *w* is the radial displacement and

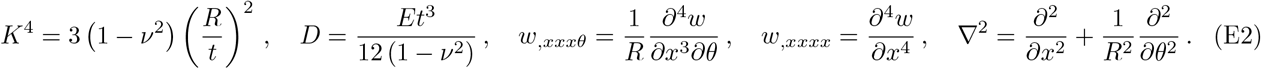

By “equal and opposite concentrated radial loads”, it is meant that two equal, spatially localized forces are applied at (*x,θ*) = (0,0) and (*x,θ*) = (0,π).

The solution to the above problem was obtained [28] by writing *q*(*x,θ*) as

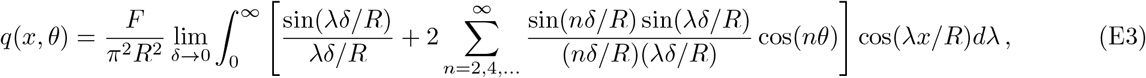

and solving Eq. (E1) order by order in *n*. The resulting deformation profile reads [28]:

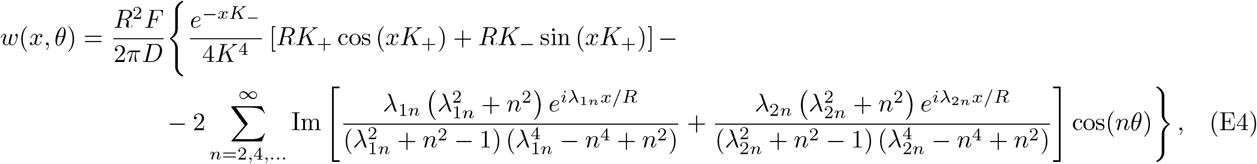

with

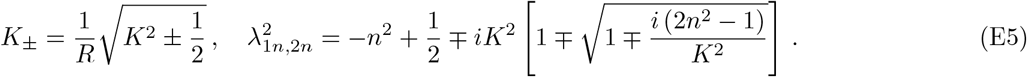

The force-indentation relation is obtained from Eq. (E4) by determining the deformation *w* at either one of the loading points, i.e. *w*(0,0) in the above formulæ, as a function of *F*. An example of the force-indentation relation and the deformation profile predicted by Eq. (E4) are shown in Fig. 1.

By using the numerical approach described in the main text, we determined the mechanical response of a thin cylindrical shell, and compared it to the above solution. Note that we are solving directly the equations of threedimensional elasticity, at variance with the simplified set of equations in [28]. Results for different mesh sizes are shown in Fig. 1: the deformation profile is indeed captured by our code, even with a relatively coarse mesh; a finer mesh is needed to capture quantitatively the force-indentation relation.

**Appendix E, Figure 1.**
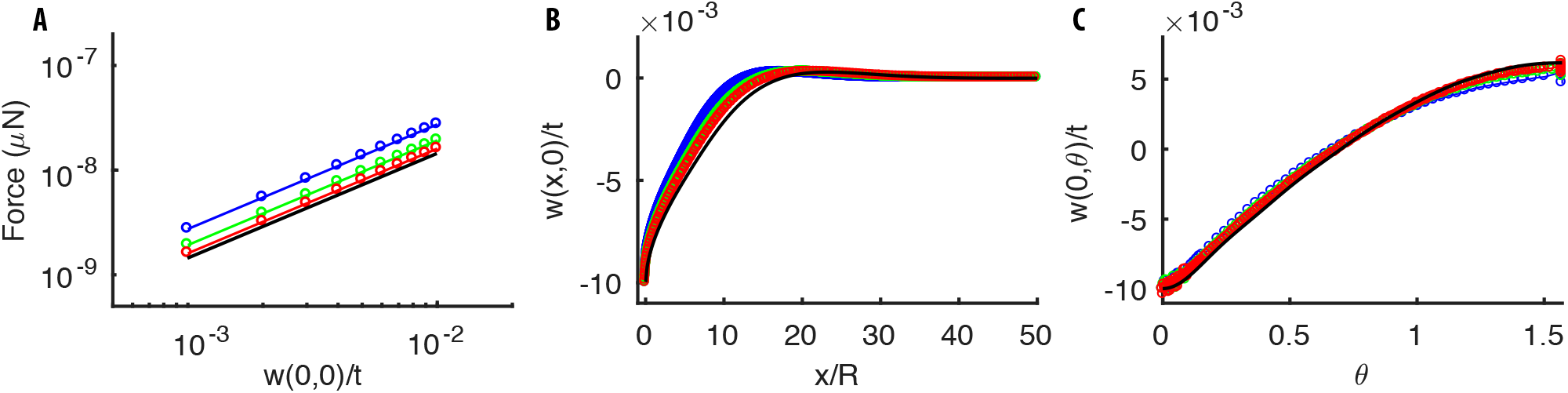
Mechanical response of a thin cylindrical shell to equal and opposite concentrated radial loads. (**A**) Force-indentation relation. (**B-C**) Radial deflection along the longitudinal and angular directions. The analytical solution Eq. (E4) (black line) is well approximated by numerical solutions (colored lines) obtained by using our numerical code. The agreement improves as the numerical mesh becomes finer (the mesh length is *t* (blue), *t*/2 (green), *t*/3 (red), where *t* is the thickness of the shell). Parameters of the simulations are: *t*/*R* = 10^−2^, *t* = 1*μ*m, *E* = 1 MPa, *ν* = 0.3.

### 2. Large indentation of spherical shell

Large indentations of pressurized spherical shallow shells were investigated in [29]. In response to a point indentation (and in the absence of buckling [56]), the 3D equations of nonlinear elasticity reduce to

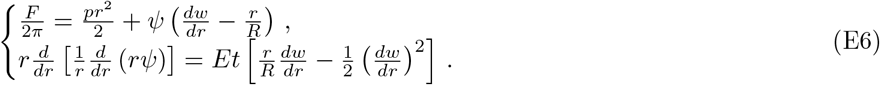

Here, *r* indicates the distance in the plane orthogonal to the indentation point, *p* is the internal pressure, *w*(*r*) is the deformation field, and *ψ* is related to the components of the stress tensor as *σ_θθ_* = *ψ*’ and *σ_rr_* = *ψ*/*r*. The nonlinear term in Eq. (E6) is due to geometrical nonlinearities generated by large deformations. Boundary conditions are:

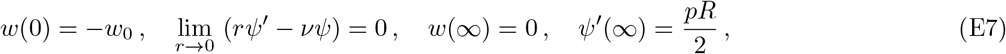

where the prime denotes differentiation with respect to *r*.

A test of our numerical scheme is provided by the comparison of its results for a thin spherical shell to the solution of the simplified Eq.(E6). Examples of force-indentation relations and deformation profiles given by Eq.(E6) for different values of *p* are shown in Fig. 2. Results of our code are also compared in Fig. 2: both the force-indentation relation and the deformation profile are well reproduced.

**Appendix E, Figure 2.**
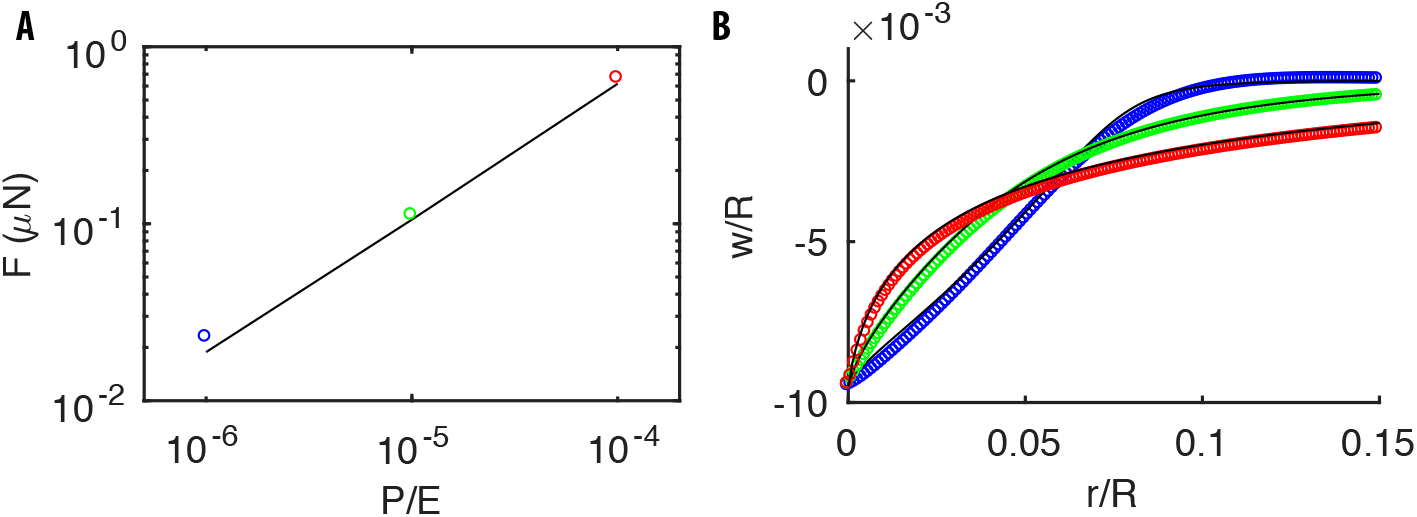
Mechanical response of a pressurized thin spherical shell to a point indentation at the north pole. (**A**) The force required to produce a deformation *w*_0_/*R* = −10^−2^ for different values of *p*. (**B**) The deformation profile for *p*/*E* = 10^−6^ (blue), 10^−5^ (green), 10^−4^ (red). Black lines and colored dots correspond to the solutions of Eq. (E6) and the results by our code, respectively. Note that, as for a pressurized cylinder (see main text), the deformation profile narrows as *p* increases. Parameters of the simulations are: *E* = 1 MPa, *ν* = 0.3, *t*/*R* = 10^−3^, *t* = 1*μ*m.

## Appendix F: How the gluing of the nematode onto the plate influences its mechanical response

Standard touch sensation experiments have the nematode glued onto a plate. As mentioned in the main text, the displacement of its body is strongly limited at locations where the glue is applied. This Appendix will analyze the effect of the gluing onto mechanical responses.

We consider the limiting case where only the line of contact with the plate is glued, i.e. the limit opposite to the one considered in the main text. There, the entire lower half of the body was glued, which was motivated by the experiments reported in the paper. In our model, the south pole of the cylinder corresponds to the line of contact with the plate in the absence of indentation. Fig. 1 shows the corresponding response of the shell to indentation.

It is of interest that the stiffness in Fig. 1A is smaller for the south-pole gluing, even though its longitudinal deformation is more extended than for the lower-half gluing, as visible in Fig. 1B. That is accounted by the deformation along the orthogonal direction, which expands in the entire lower half of the body (see Fig. 1C). That deformation is forbidden for the lower-half gluing, which is the reason for the increased stiffness. Generally, the stiffness is expected to decrease as we reduce the region where the nematode is glued to the plate, which could be verified experimentally.

**Appendix F, Figure 1.**
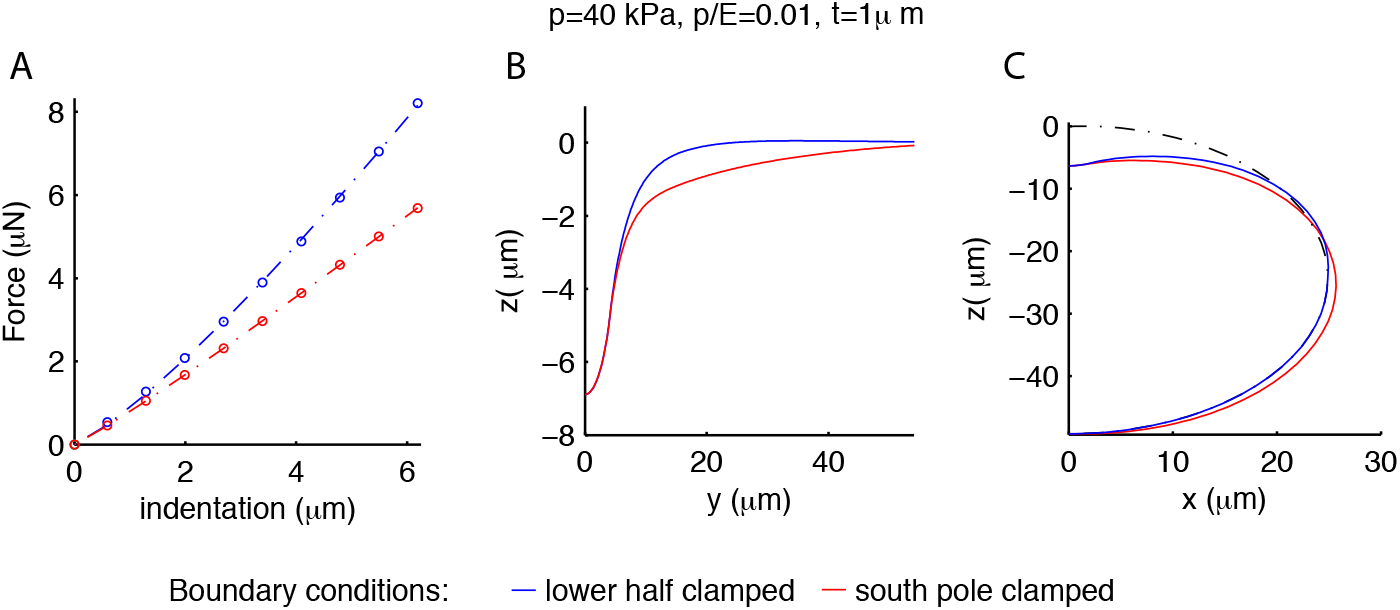
Influence of the gluing of the worm on its mechanical response. Blue and red curves correspond respectively to gluing of the entire lower half of the cylinder or the line of contact with the plate (south pole) only. (**A**) Force-indentation relations; note that the stiffness is greater for the blue curve. The deformation profiles along the longitudinal (**B**) and orthogonal (**C**) directions are wider for the south-pole gluing, which has the lower half of the shell deformed in the orthogonal direction as well. The undeformed geometry is the black dash-dotted line.

## Appendix G: Influence of the boundary conditions at the ends of the cylinder on its mechanical response

The main text discusses a pressurized cylindrical shell with free conditions at the two ends of the cylinder. Here, we analyze how mechanical properties are modified when the ends are closed by plugs, which leads to an additional component of stress.

We computed numerically the response to the indentation of a closed pressurized cylindrical shell. The numerical procedure is quite analogous to the main text, with the only difference of the boundary conditions. The action of the pressure on the plugs produces a longitudinal force on the shell, whose magnitude does not depend on their structure (but for a boundary layer close to the ends). Without loss of generality, we used semi-spherical plugs.

Results of the simulations are shown in Fig. 1. The deformation profile for a given value of *p*/*E* is more extended than for free conditions at the sides (Fig. 1A). Since the associated change in volume is bigger, and the stiffness is dominated by internal pressure, the stiffness of the shell is greater (Fig. 1B). The additional stiffness stems from the longitudinal stress introduced by the lateral plugs, as confirmed by the decrease of the difference between the closed and the open conditions with the internal pressure (Fig. 1A,B).

Let us now show that closed ends cannot reproduce experimental data, which is the reason why the main text focuses on open cylinders. We first fix the range *p*/*E* ∈ [0, 0.02] considered so far. The extension of the deformation decreases with *p*/*E* but, even at the largest value *p*/*E* = 0.02, it remains too wide to account for experimental data. Further increase in *p*/*E* does reduce the extension, yet it runs in conflict with experimental data on the bulk modulus [37]. Indeed, the estimate for *p* obtained in the main text depends only on the deformation profile and the force-indentation relation, which are both given by the experiments. We can therefore fix *p* = 40kPa, and predict the bulk modulus for the corresponding various values of *E*. Results in Fig. 1C are systematically smaller than experiments (and even further increase of *p*/*E* would not help as the bulk modulus decreases with *p*/*E*).

In summary, we showed that the description of the nematode body as an elastic shell with closed ends cannot reproduce experimental data on the mechanics of *C. elegans*. Conversely, the main text showed that the same elastic model with free sides does capture main features of the mechanical response. Our results suggest that the longitudinal stress generated by the plugs is somehow relaxed in the worm, which may relate to the annular structure of the cuticle.

## Appendix H: The local geometry of the channels along the TRNs

We treat the TRN as a (small) cylinder running (at rest) in the upper part of the (big) cylinder in Fig. 1, at *x* = 0 along the *y* axis. The TRN is placed right above the internal part of the shell. Upon indentation, the orthogonal basis x̂_i_ (*i* = 1,2,3) in Fig. 1 is deformed into a triad of vectors 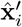 in a way that depends on its original location. The separation between pairs of neighboring material points (**r,r**+**dr**) is calculated using the gradients of the displacement field **u**(**r**). In particular, the variation of the squared distance *dr’*^2^ − *dr*^2^ = 2*ε_ij_ dr_i_ dr_j_*, and the angle between two vectors **dr_1_** and **dr_2_** (see [20, 22])

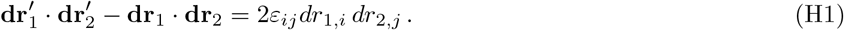

**Appendix G, Figure 1.**
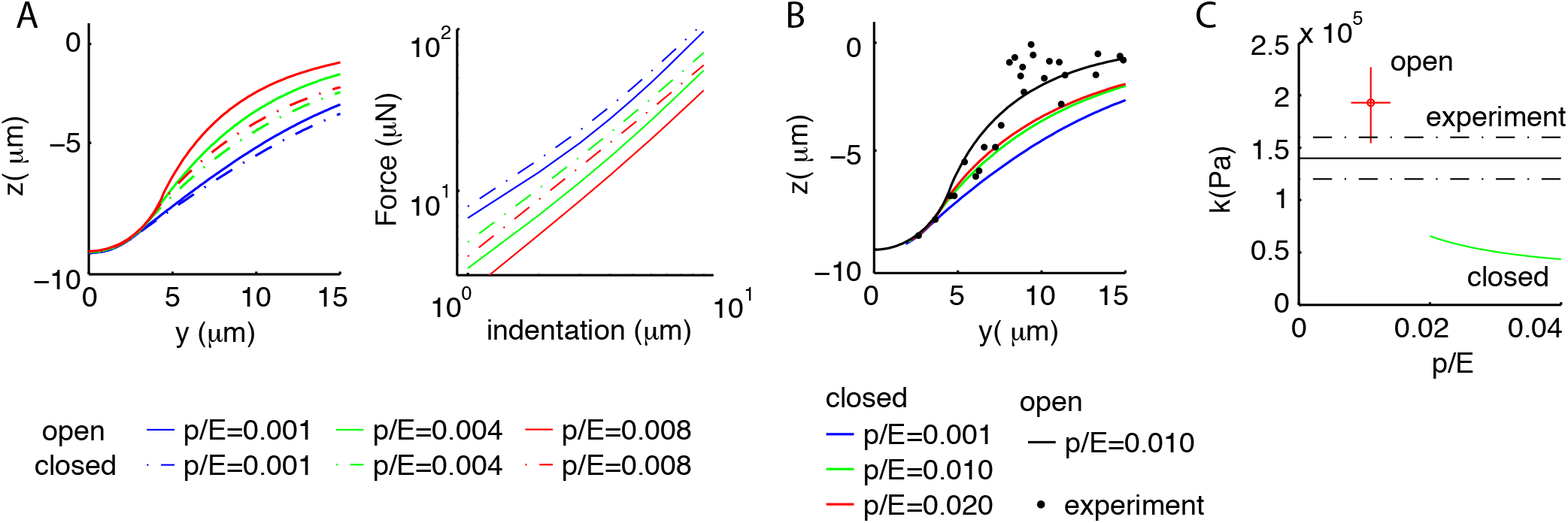
Boundary conditions at the ends of the shell influence mechanical properties. (**A**) Comparison of the mechanical response for an open (dash-dot line) and closed (continuous line) cylinder. For a given value of *p*/*E*, the deformation profile is more extended (left) and the shell stiffer (right) if the two ends are closed; the difference increases with *p*/*E*. (**B**) Experimental and numerical deformation profiles along the longitudinal axis. None of the values *p*/*E* ∈ [0, 0.02] for the closed cylinder (colored line) captures experimental data. (**C**) Experimental and predicted values for the bulk modulus. The value predicted for closed conditions at the ends decreases with *p*/*E* and is too small to account for the data. Results for free lateral conditions are shown for comparison. Parameters of the simulations are as in Fig. 2 of the main text.

A convenient orthonormal basis 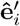 to analyze the dynamics of the channels is defined as follows: 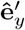 is aligned with the local direction of the (deformed) axis of the cylinder running head-to-tail; 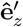 is orthogonal to the neural membrane at the top of the TRN, and oriented outward; 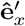 is tangential to the neural membrane, along the remaining direction of a right-handed system.

For every channel, we define a local base **ŵ**_*i*_ such that **ŵ** _1,2_ span the plane locally tangential to the neural membrane while **ŵ**_3_ indicates the orthogonal direction. The bases **ŵ**_i_ are constructed by rotating the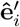 appropriately. For a channel placed at the top of the TRN, the local basis **ŵ**_i_ coincides with 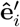. If the channel is rotated by *θ* along the surface of the TRN, then 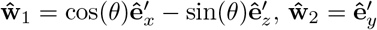, and 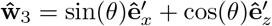.

## Appendix I: Comparison of different functional forms for the activation of the channels

In our model, the free energy of a channel is modulated by the deformation of its elastic filament. A general rotationally-symmetric form of the free energy reads

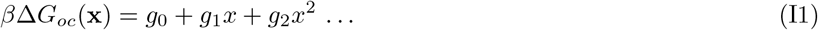

where *x* = |**x**|. In the main text, we used the linear form of Eq. (I1); here, we repeat the analysis of the experimental data for *g*_1_ = 0, i.e. a quadratic form. Results are shown in Fig. 1, with results for the linear model included for the sake of comparison. The quadratic dependence limits the sensitivity range, and leads to a worse description of the data (see Fig. 1); the same also holds for individual realizations of the responses (data not shown). That is witnessed by the step response in Fig. 1, where the current predicted by the model saturates at smaller values than the peaks observed in the data, even though the best-fitted baseline activity is stronger.

We also repeated the analysis of the directional model replacing Eq. (13) of the main text by

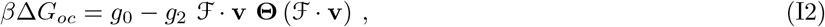

where Θ is the Heaviside function. Results were comparable to those presented in the main text.

## Appendix J: Noise analysis for non-equally stimulated channels

Here, we compute mean and variance of the current as a function of the statistics of the ion channels. We use these results in the main text to compare model predictions to experimental data. The point is to generalize standard noise analysis to the case where the channels are not equivalent. That can be the case either because they are not identical or, as it the case here, because their stimulation differs due to their location with respect to the stimulation. We also calculate the scaling of the level of fluctuations expected in a finite sample of *N* measurements.

**Appendix I, Figure 1.**
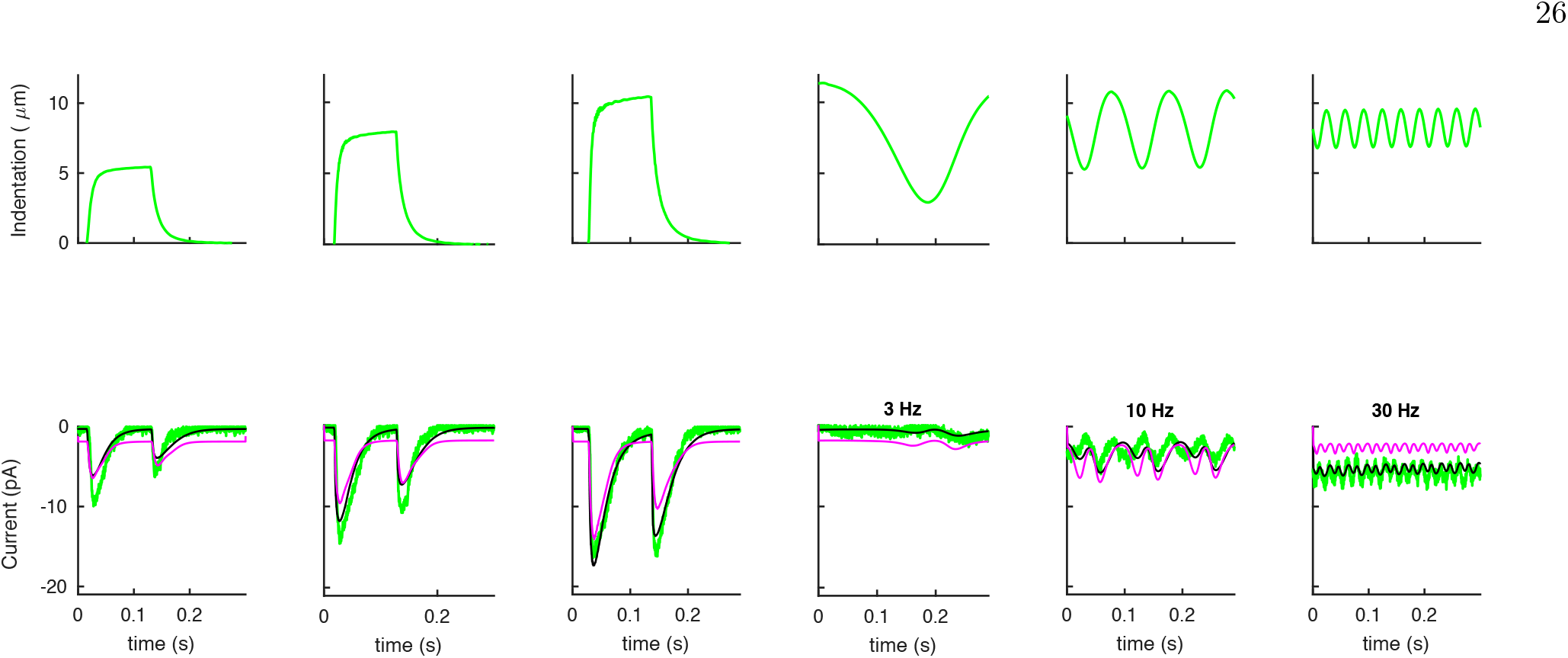
The linear form of Eq. (I1) outperforms its quadratic counterpart in describing experimental data. (A) The average predictions for the experimental data (green) given by the linear (black) and quadratic (purple) forms of Eq. (I1).

We start by deriving the expression of the mean and variance of the neural current from the statistics of the single channels. The total ion current *I* through the neuron is the sum 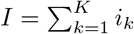, where *i_k_* is the current flowing through the *k*-th channel and the sum runs over the channels along the neuron. Channels can take three distinct states in our model: open, with maximal current *i*_0_ and probability *P_o_*(*k*); sub-conducting, with intermediate current *i_s_* and probability *P_s_*(*k*); closed, with no current and probability *p_C_*(*k*) = 1 – *p_o_*(*k*) – *p_s_*(*k*). Each channel follows a generalized Bernoulli distribution: the associated mean and variance are

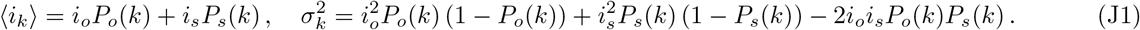

Assuming that the gatings of the channels are independent random variables, we obtain

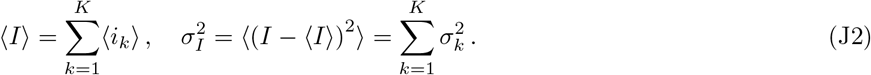

In the main text we suggested that the variance of the current could be used to study experimentally the microscopic properties of ionic channels. Since the variance should be inferred from a finite sample, we want to quantify the scaling of fluctuations in the sample variance with the number of measurements. Given *N* measurements *I_n_* of the TRN current, the sample mean *m*_1_ and sample variance *m*_2_ are defined as

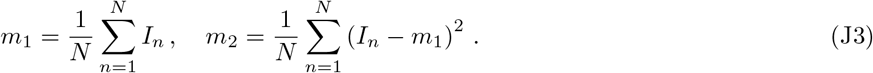

The expected sample variance and its variance are [33, 34]:

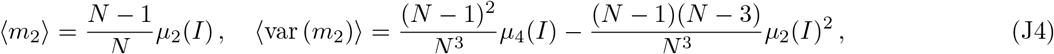

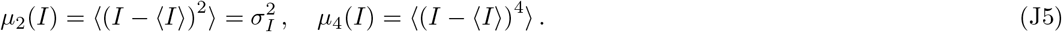

To determine the relation between the moments *μ*_2_(*I*), *μ*_4_(*I*), … of the total current *I* and the statistics of the single-channel currents, it is convenient to use cumulants and the cumulant-generating function of *I*, defined as *Q_I_*(*t*) = log〈exp(*tI*)〉 [35]. Its advantage is that the function is additive:

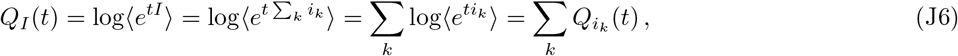

where we have exploited independence among the *i_k_*’s. Since the cumulant *q_n_*(*I*) of order *n* (see [35]) is 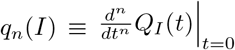, Eq. (J6) implies the additivity *q_n_*(*I*) = Σ*_k_ q_n_*(*i_k_*) of the cumulants.

The cumulants *q_n_*(*i_k_*) are calculated using the fact that the channels obey a generalized Bernoulli distribution:

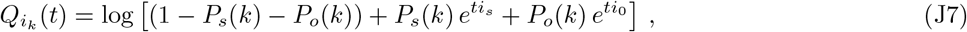

whence we obtain

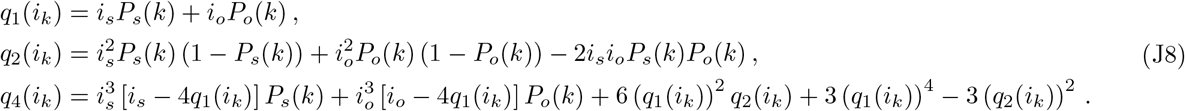

The derivation is completed by relating the central moments *μ*(*I*) to its cumulants via standard formulæ[35], e.g.

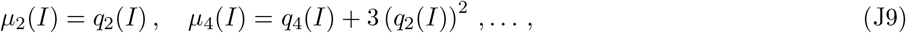

by expressing *q_n_*(*I*) as Σ*_k_ q_n_*(*i_k_*), and finally using Eqs. (J8).

## Appendix K: Mechanical effects of mutations in proteins of the cuticle

Mutations of proteins composing the cuticle have been used as a probe to investigate mechanical properties of *C. elegans* [24, 37]. This Appendix will explore the consequences of the simplest possible assumptions on the effects of those mutations, obtain predictions for our mechanical model, and compare them to experiments.

We shall describe the effects of mutations in the cuticle through variations in the stretching stiffness *S*, which is the only parameter related to the thin external layers (see Appendix D). We assume that mutations do not affect the internal pressure of the nematode. The argument in [24] is that the genes involved, e.g. *dpy-5* and *lon-2*, do not affect transport proteins likely to regulate osmotic pressure. It is possible, though, that mutations in the cuticle affect the development of the nematode, hence its body structure and the internal pressure. To constructively advance the issue, we assume that that does not happen and explore consequences.

Fig. 1A shows the dependence of geometrical properties on *S*: the radius *R* of the pressurized cylinder decreases, whilst its length *L* increases with *S*. This reflects changes with respect to the unpressurized condition brought by the internal pressure *p*. Fig. 1B shows that the bulk modulus increases with *S*. Finally, the stiffness *S* enters the response to indentation experiments. Fig. 1B shows the ratio *f* ≡ *F*/*w*_0_, i.e. the force *F* needed to reach an indentation *w*_0_ = 5*μm*, *υs S* (we verified that results do not depend on the choice of *w*_0_). The increase with *S* is due to two contributions that affect the change in volume via the deformation field. First, if the external radius of the shell is kept constant, the deformation field is wider (see Fig. 2B). Second, the radius of the shell becomes smaller as *S* increases (see Fig. 1A) which leads to a larger deformation field (data not shown).

The parameter *S* is not measured experimentally, which forces us to use *R* as a proxy. In this formulation, the model predicts that mutations in the cuticle which increase *R* will decrease *L*, stiffness and bulk modulus. These features are in qualitative agreement with experimental observations [24, 37].

Quantitatively, we observe that: (i) the length *L* of the mutants in [24] is systematically larger than our predictions; (ii) the radius of *lon*-2 mutants in [24], which is about 25% smaller than the wild type, cannot be obtained in our simulations, as seen in Fig. 1C. (i) may be due to the likely non-isotropy of the Young’s modulus generated by the annular structure of the external layers. That is expected to make the stiffness smaller in the longitudinal than the orthogonal direction. (ii) is due to the fact that *R* cannot be smaller than the value for the unpressurized shell, which is only 20% smaller than the wild type. Since relative changes in *R* grow with *p*/*E*, it is likely that uncertainties on *R* are due to fluctuations in **p*/*E** among worms (see main text). As already mentioned above, both *R* and *L* could also be affected by our neglecting developmental effects.

We finally compare in Fig. 1C-D our predictions to experimental data from [24, 37]. The agreement is notable and leads to the prediction that the bulk modulus in *lon*-2 mutants should significantly deviate from the wild type. This differs from the conclusion that the bulk modulus is not affected by the cuticle [37], which was based on mutants other than *lon*-2.

**Appendix K, Figure 1.**
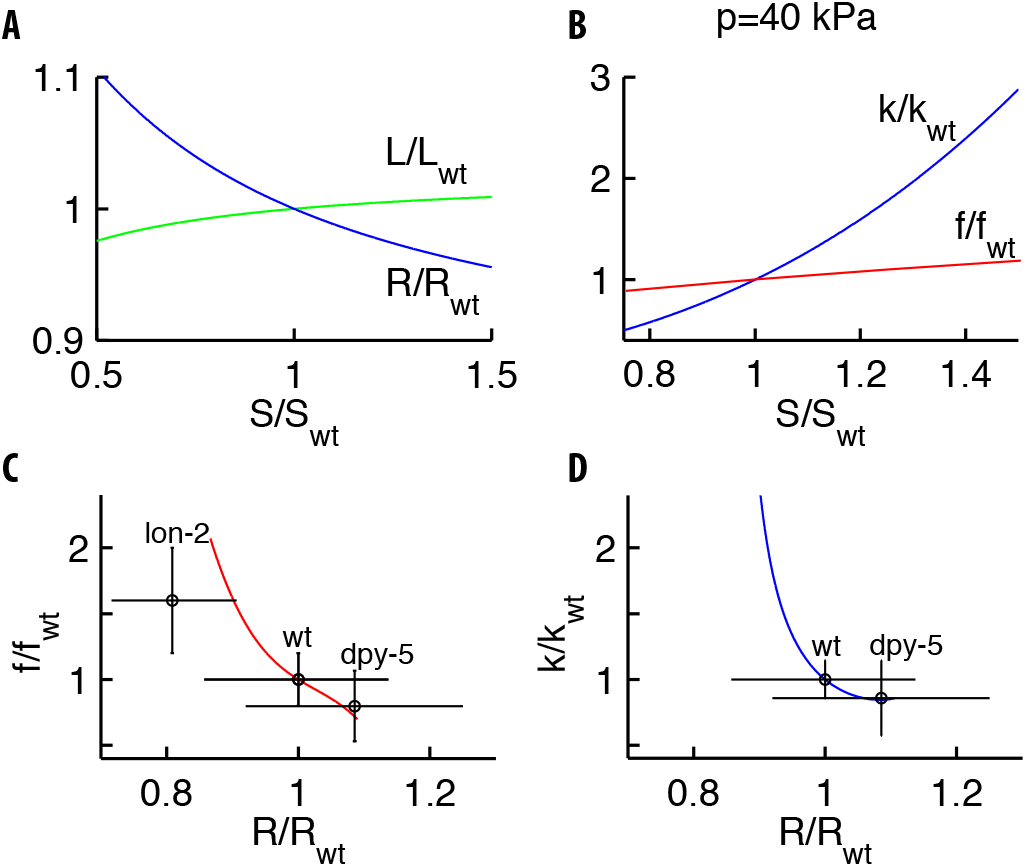
Changes in the mechanics caused by mutations of proteins in the cuticle. Mutations are modeled via changes of the stretching stiffness *S* with respect to the wild type *S_wt_*. (**A**) As *S* increases, the geometry of the pressurized cylinder modifies: its length *L* increases and its radius *R* decreases. (**B**) As S increases, the bulk modulus *k* (blue) and the stiffness *f* (red) of the force-indentation relation increase. (**C-D**) Our theoretical predictions and experimental measurements [24, 37] for *f* and *k υs R*. Since the radius of the mutants was not reported in [37], we used values in [24]. The upshot is that *lon* −2 mutants should have a bulk modulus significantly different than the wild type.

## Appendix L: Inference of the model parameters

The amplitude of **Γ**(0), which controls the scale of the elongation in Eq. (5), is set to unity by redefining the parameters *g_h_* and *g_s_* in Eq. (8). As for the rates *R*, a parsimonious form that respects Eqs. (7) and (10) is

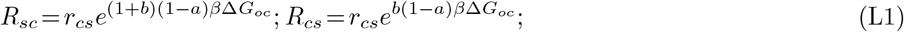

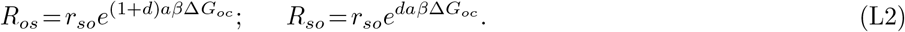

Here, *r_cs_* (*r_so_*) controls the rate of the transitions between the closed and the subconductance states (the subconductance and the open states) and the parameters *b, d* ∈ [−1, 0] control their global shift with respect to variations of the free-energy difference. Parameters are inferred from experimental curves by using the MATLAB optimization function “lsqcurvefit”, based on the least-squares distance between the predicted and the observed current profiles. For every realization of the quenched disorders (distribution of the channels and the initial direction of their filaments), we obtain the best parameters, which are then averaged. Their means are *τ* = 1.4ms; *g_h_* = 1.410^−3^; *g_s_*/*g_h_* = 0.09; *r_cs_* = 1/69.5ms; *b* = −0.75; *r_so_* = 1/18ms; *d* = −0.56; a = 0.50; *i_s_*/*i*_0_ = 0.71.

## Appendix M: Dependence of neural responses on the properties of elastic filaments

This Appendix will investigate the dependence of the TRN neural response on the interaction between the channels and the surrounding medium, which we have embodied here into an elastic filament. In our model, those interactions are described by two parameters: the elastic constant *k* of the filament, and its friction coefficient *γ* with the medium. The neural response depends on the ratio *τ* = *γ*/*k*, which appears in Eq. (5) of the main text.

To understand the effects of *τ*, we computed the TRN current predicted by our model in response to a step of fixed amplitude as a function of *τ*, keeping fixed all other parameters detailed in the main text. Fig. 1 shows the peak current and the decay time, i.e. the time for the current to reach half of its peak value, both averaged over the statistical realizations of the distributions for the channels. Both the peak current and the decay time increase with *τ*: filaments relax more slowly, which provides a stimulus on the associated channel that lasts longer and thereby allows for a higher current.

Fig. 2 shows the histogram of least-squares errors for individual responses to the stimuli in Fig. 3 of the main text for different geometries of the filaments attached to the channels. The black curve shows that individual realizations as well (and not just the average as in the main text) reproduce neural responses for the various profiles, strengths and frequencies. The figure also presents results for a model with filaments initially or permanently restricted to be tangential. Graphs indicate that our predictions are further improved by introducing those additional assumptions.

**Appendix M, Figure 1.**
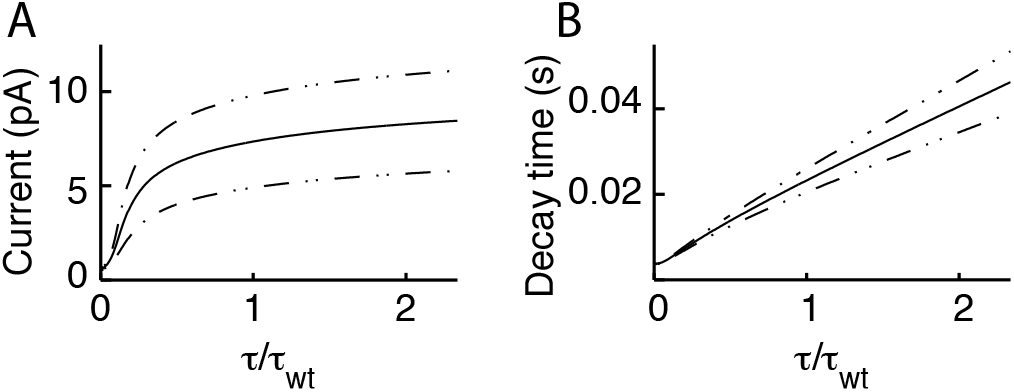
Dependence of the neural response on the filament-medium interaction. Peak current (**A**) and decay time (**B**) as a function of the relaxation time *τ* of the elastic filament connected to the channel. The model predicts that both the peak current and the decay time increase with *τ*.

**Appendix M, Figure 2.**
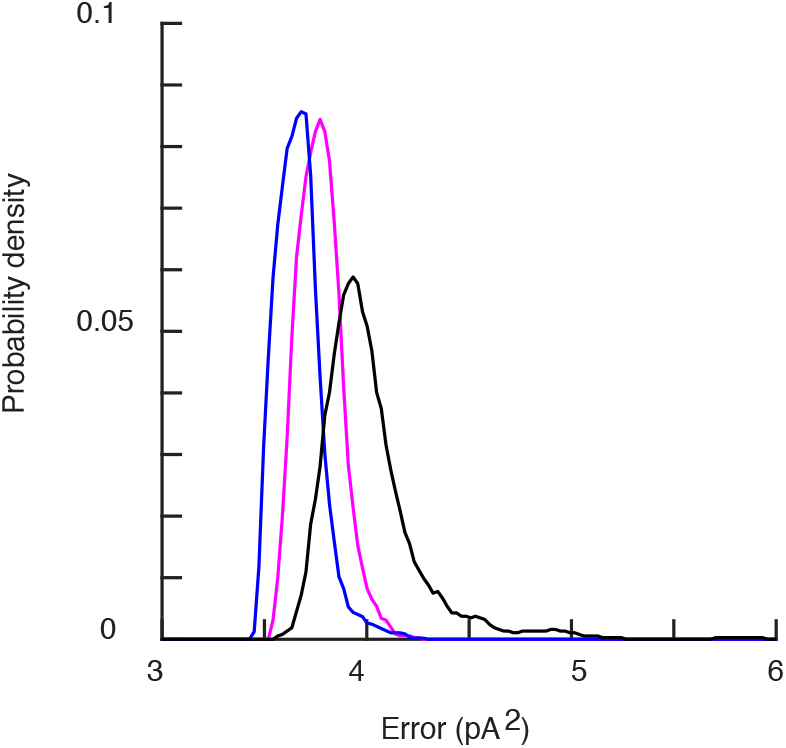
Dependence of the neural response on the geometry of the filaments. The error for the profiles of stimulation in Fig. 3 of the main text. The black histogram is built from individual realizations for the unconstrained model discussed in the main text. The purple and the blue curves refer to the corresponding histograms for filaments initially or permanently restricted to be tangential to the neural membrane.

